# *S. pombe* Mto1[bonsai]:Mto2 stabilizes γ-tubulin positions within the γ-TuRC

**DOI:** 10.64898/2026.11.06.764827

**Authors:** James D. Le Cornu, Su Ling Leong, Eric M. Lynch, Harish C. Thakur, Kenneth E. Sawin, Atlanta G. Cook

## Abstract

Microtubules are present in all eukaryotes and are nucleated by multiprotein γ-tubulin ring complexes (γ-TuRCs), ring-shaped structures that contain several copies of the γ-tubulin small complex (γ-TuSC). γ-TuRCs nucleate microtubules via a template-based mechanism in which the positions of γ-tubulins in the γ-TuSCs match the position of αβ-tubulin dimers in the microtubule lattice. Although γ-TuSC proteins are highly conserved, other components of the γ-TuRC can vary in different organisms, as can their relative importance. Here, we characterize two different simplified γ-TuRCs from fission yeast Schizosaccharomyces pombe, using recombinant purified proteins and negative stain electron microscopy. We show that a nucleation-incompetent γ-TuRC that contains only γ-TuSC proteins plus the microprotein Mzt1 (γ-TuSC:Mzt1) can form rings as efficiently as a large, nucleation-competent complex (MGM) that additionally contains the proteins Mto1 and Mto2. We find that the position of γ-tubulins in γ-TuSC:Mzt1 are less well defined than in MGM. Comparative modelling using our electron microscopy and crosslinking mass spectrometry data, together with structures of γ-TuRCs from other organisms, suggests that Mto1 and Mto2 form a multimeric outer mantle surrounding the γ-TuSC:Mzt1 γ-TuRC. This mantle may stabilize positions of γ-tubulins within MGM, explaining the microtubule-nucleating ability of MGM and highlighting the diversity of non-γ-TuSC proteins in modulating γ-TuRC structure and function.

## Introduction

Gamma-tubulin ring complexes (γ-TuRCs) are multimeric, megadalton-sized protein complexes important for the efficient nucleation of microtubules (MTs) in vivo (Moritz et al, 2000; Kollman et al, 2011; Roostalu & Surrey, 2017). γ-TuRCs allow MT nucleation at concentrations of αβ-tubulin dimer lower than is required for spontaneous nucleation by tubulin dimer alone (Johnson & Borisy, 1977; Evans et al, 1985a; Job et al, 2003; Thawani et al, 2020). Structural studies of γ-TuRCs in the absence of growing MTs reveal a low-pitch conical structure with a lock-washer geometry at the base that is similar, but not identical to the geometry of MTs (Oegema et al, 1999; Gunawardane et al, 2000a; Kollman et al, 2010). Structures of γ-TuRCs with nucleated MTs show a shift to a closed lock-washer state that matches the helical pitch of MTs and accommodates precisely 13 protofilaments (Aher et al, 2024; Brito et al, 2024; Dendooven et al, 2024). In contrast, MTs polymerized from tubulin dimer alone typically contain 12-16 protofilaments (Evans et al, 1985b). An emerging picture is that structural rearrangements within the γ-TuRCs, perhaps cooperatively with the templated MT protofilaments, result in the rearrangement of the γ-tubulins and GCPs to align with MT geometry (Brito et al, 2024).

Although γ-TuRCs from different organisms share a common lock-washer architecture, they also have significant compositional and structural differences (Kollman et al, 2011; Roostalu & Surrey, 2017; Tovey & Conduit, 2018; Sulimenko et al, 2022). All γ-TuRCs contain multiple copies of γ-tubulin small complexes (γ-TuSCs). The γ-TuSC is a Y-shaped heterotetramer containing two γ-tubulins, one γ-tubulin complex protein 2 (GCP2) and one γ-tubulin complex protein 3 (GCP3); GCP2 and GCP3 are paralogs, and each binds one γ-tubulin (Suppl. Fig. 1A). In addition, many eukaryotic γ-TuRCs contain the proteins GCP4, GCP5, and GCP6, which are similar in sequence to GCP2 and GCP3. GCP4 can pair with GCP5 and with GCP6 to form (together with γ-tubulins) γ-TuSC-like complexes (Suppl. Fig. 1B). Structural analyses have revealed that in H. sapiens and X. laevis γ-TuRCs, four γ-TuSCs are followed by a GCP4-GCP5 γ-TuSC-like complex, a GCP4-GCP6 γ-TuSC-like complex, and a γ-TuSC at the lock-washer seam (Consolati et al, 2020; Liu et al, 2020; Wieczorek et al, 2020b; Zimmermann et al, 2020; Würtz et al, 2022; Aher et al, 2024; Dendooven et al, 2024; Gao et al, 2025) (Suppl. Fig. 1C). The two γ-TuSC-like complexes form a salt-stable subcomplex that is thought to promote assembly of the γ-TuRC (Fava et al, 1999; Oegema et al, 1999; Gunawardane et al, 2000a; Murphy et al, 2001; Anders et al, 2006; Vérollet et al, 2006; Choi et al, 2010; Consolati et al, 2020; Haren et al, 2020; Liu et al, 2020; Wieczorek et al, 2020b). Interestingly, while GCP4, GCP5, and GCP6 are present in all eukaryotic kingdoms, they appear to have been lost (or are extremely diverged) in certain lineages (Haruta et al, 2023). For example, although most fungi have GCP4, GCP5, and GCP6, they are largely absent from the Saccharomycotina, a subphylum that includes budding yeast Saccharomyces cerevisiae (Suppl. Fig. 1D). GCP4, GCP5, and GCP6 are also absent—or at least not identifiable—in a Nematode lineage that includes Caenorhabditis elegans (Haruta et al, 2023). Moreover, in at least some organisms that do have GCP4, GCP5 and GCP6, these proteins are not strictly required for MT nucleation (Anders et al, 2006; Vogt et al, 2006; Tovey & Conduit, 2018). Another important difference among γ-TuRCs from different eukaryotes involves the microprotein Mzt1 (MOZART1 in H. sapiens and MZT-1 in C. elegans) (Dhani et al, 2013; Masuda et al, 2013a, 2013b; Cota et al, 2016; Lin et al, 2016b; Tovey et al, 2018; Leong et al, 2019; Huang et al, 2020). MOZART1 and Mzt1 are essential in H. sapiens and fission yeast Schizosaccharomyces pombe, respectively, but the ortholog is non-essential in D. melanogaster and absent from S. cerevisiae altogether (Seltzer et al, 2007; Nakamura et al, 2012; Dhani et al, 2013; Masuda et al, 2013a; Lin et al, 2016b; Tovey et al, 2018). In vertebrate γ-TuRCs, Mzt1 forms complexes with the N-terminal extensions (NTEs) of GCP3 and GCP5 through conserved interactions (Huang et al., 2020). Overall, this shows that different organisms have evolved different approaches to solving the “MT nucleation problem” (Tovey & Conduit, 2018; Gao et al, 2024).

In S. pombe, GCP4, GCP5 and GCP6 are all present in the γ-TuRC and enhance MT nucleation efficiency in vivo, but they are not essential for nucleation (Murphy et al, 2001; Fujita et al, 2002; Venkatram et al, 2004; Anders et al, 2006; Masuda et al, 2013a; Yukawa et al, 2019). Additional factors critical for MT nucleation in vivo in S. pombe are the proteins Mto1 and Mto2. Mto1 contains a Centrosomin Motif 1 (CM1) domain, a conserved sequence also found in S. cerevisiae spindle pole body component Spc110 and vertebrate CDK5RAP2. CM1-domain proteins interact with GCP2, promoting a γ-TuRC lock-washer geometry closer to MT geometry (Wieczorek et al, 2020a, 2020b; Brilot et al, 2021; Serna et al, 2024, 2026; Xu et al, 2024). Mto2, which is found in many fungi, binds to Mto1 and is thought to form higher-order complexes with Mto1 (Sawin et al, 2004; Janson et al, 2005; Samejima et al, 2005; Venkatram et al, 2005; Lynch et al, 2014; Leong et al, 2019; Thakur et al, 2019).

Previously we showed that S. pombe γ-TuSCs (containing γ-tubulins, GCP2, and GCP3) can be reconstituted into larger assemblies from purified recombinant proteins, provided that Mzt1 is present to stabilise recombinant GCP3 (Leong et al, 2019). However, these assemblies (“γ-TuSC:Mzt1 complexes”) do not promote MT nucleation in vitro. Reconstitution of nucleation-competent assemblies requires the addition of Mto1[bonsai] (a CM1 domain-containing fragment of Mto1) and Mto2 (Leong et al, 2019). We refer to the nucleation-competent assembly as the “MGM complex”, as it contains: <u>m</u>icrotubule organizer proteins Mto1 and Mto2; <u>γ</u>-TuSC proteins γ-tubulin, GCP2, and GCP3; and <u>M</u>zt1 (Suppl. Fig. 1E). The MGM complex, a simplified system relative to what may exist in vivo in S. pombe, appears to represent a minimal complex sufficient for MT nucleation in vitro (Lynch et al, 2014; Leong et al, 2019).

While the overall architecture of γ-TuRCs and mechanisms of templated MT nucleation are likely to be similar across species, it is currently not known how Mto1 and Mto2 convert γ-TuSC:Mzt1 into a nucleation-competent assembly, nor how the MGM structure compares to well-characterized vertebrate γ-TuRCs. Previously we showed by negative stain electron microscopy (NSEM) that MGM complexes form rings (Leong et al, 2019). Here we use NSEM to characterize and compare MGM and γ-TuSC:Mzt1 in greater detail. We show that γ-TuSC:Mtz1 complexes also form rings, with a similar efficiency to that of MGM complexes. In addition, through 3D reconstruction, we find that both complexes form conical lock-washer structures characteristic of γ-TuRCs, but with differing geometries. Our data suggest that rather than assisting γ-TuSC:Mzt1 complexes to form rings, Mto1[bonsai] and Mto2 reduce conformational heterogeneity in MGMs, promoting a suitable template for MT nucleation. Finally, by combining AlphaFold3 models with prior crosslinking mass spectrometry data, we suggest how Mto2 interacts with Mto1 to form Mto1:Mto2 complexes, and how Mto1:Mto2 complexes may form an outer layer on the γ-TuRC lock-washer.

## Results

### γ-TuSC assemblies form ring structures *in vitro*

We purified γ-TuSC:Mzt1 and MGM complexes from baculovirus-mediated expression as previously described (Leong et al, 2019). The expected stoichiometric ratio of the γ-TuSC:Mzt1 complex components is 2:1:1:1 γ-tubulin:GCP2:GCP3:Mzt1, while the expected ratio of components in the MGM complex is ∼ 2:2:2:1:1:1 Mto1[bonsai]:Mto2:γ-tubulin:GCP2:GCP3:Mzt1 (Oegema et al, 1999; Lynch et al, 2014; Greenberg et al, 2016; Leong et al, 2019). To estimate the relative stoichiometries of proteins in each recombinant complex, dilution series of MGM and γ-TuSC:Mzt1 were analysed using SDS-PAGE (Fig. 1A). Mto2 appears as a disperse band, due to phosphorylation at multiple sites (Borek et al, 2015). Mzt1 migrated at the dye front and was not quantified. Based on the integrated intensity of the bands and molecular weights, we estimated stoichiometries to be 2:1:1 γ-tubulin:GCP2:GCP3 for γ-TuSC:Mzt1, and 2:2:3:1:1 for Mto1[bonsai]:Mto2:γ-tubulin:GCP2:GCP3 for the MGM complex. The γ-tubulin stoichiometry in the MGM complex is likely overestimated, as Mto2 and γ-tubulin bands overlap. Overall, this analysis suggests that within MGM, there are two molecules of Mto1[bonsai] and two of Mto2 for each GCP2-GCP3 pair, consistent with previous quantitation in vivo (Lynch et al, 2014).

**Figure 1:**
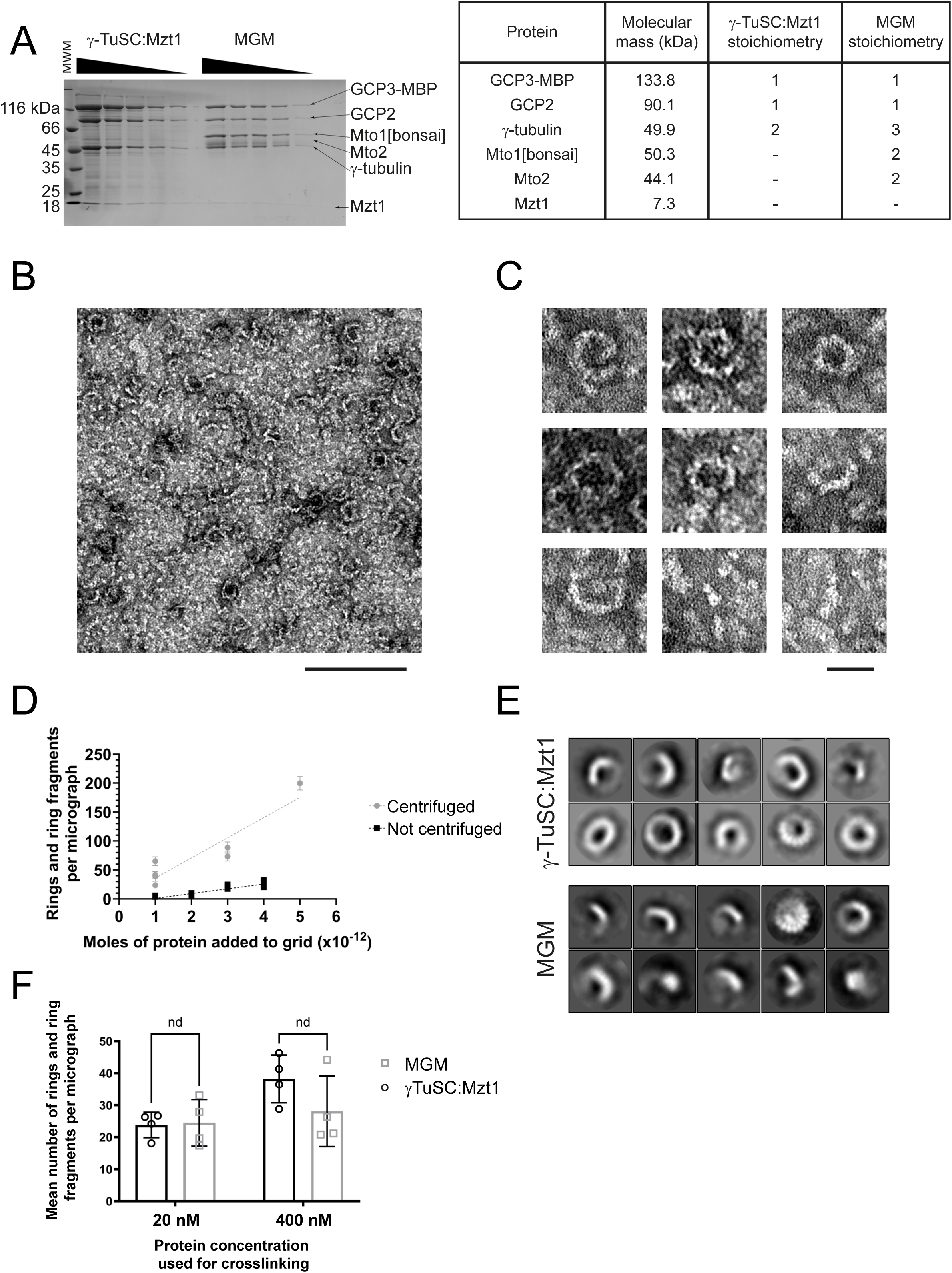
γ-TuSC:Mzt1 forms ring-shaped and partial ring-shaped complexes at a similar efficiency to MGM complex. **A)** Quantitative analysis of γ-TuSC:Mzt1 and MGM by Coomassie-stained SDS-PAGE, with a table of calculated subunit stoichiometries. **B)** Negative stain electron micrograph of γ-TuSC:Mzt1 (130 nM concentration). **C)** Selected representative particles of γ-TuSC:Mzt1 from multiple micrographs as in B. Shapes observed: ring, spiral, crescent and rod. **D)** Quantification of number of γ-TuSC:Mzt1 ring and ring-fragment complexes. Each symbol indicates mean number of rings and ring fragments from a dataset collected from a single grid. Error bars indicate standard deviation for each grid. **E)** Examples of most populated ring and ring fragment-shaped 2D class averages of γ-TuSC:Mzt1 and MGM. **F)** Quantitative negative stain electron microscopy (NSEM) assay comparing the number of ring and ring-fragment particles per micrograph, for γ-TuSC:Mzt1 and MGM, each crosslinked with glutaraldehyde at the indicated protein concentrations prior to dilution and analysis by NSEM. Each hollow circle or square indicates the mean number of ring and ring-fragment particles from a single grid. The column indicates the mean of mean number of ring and ring-fragment particles. Error bars represent the standard deviation of the mean number of ring and ring-fragments of each grid. Scale bars: (B) 150 nm; (C) 30 nm.

Previous glycerol gradient centrifugation experiments suggested that γ-TuSC:Mzt1 forms oligomers containing multiple copies of γ-TuSCs (Oegema et al, 1999; Gunawardane et al, 2000b; Vinh et al, 2002; Leong et al, 2019), but whether these oligomers are rings was not investigated. Using NSEM we visualized γ-TuSC:Mzt1 particles at concentrations ranging from 40-130 nM (∼0.1-0.3 mg/mL). We observed particles of different sizes and shapes, including rings, spirals, crescents, and rods, with lengths ranging from 10-35 nm (Fig. 1B, C). The presence of rings and spirals, which likely consist of multiple copies of γ-TuSC:Mzt1, indicates that neither Mto1 nor Mto2 is required for ring formation. The presence of crescents, which may be partial rings, further suggests that the oligomeric state of γ-TuSC:Mzt1 can vary. Rods likely correspond to single hetero-pentamers of γ-TuSC:Mzt1.

Given that both MGM and γ-TuSC:Mzt1 have the ability to form rings, but only MGM efficiently nucleates microtubules in vitro, we hypothesized that Mto1[bonsai]:Mto2 might facilitate the efficiency of ring formation (Kollman et al, 2010, 2011; Lynch et al, 2014; Farache et al, 2016, 2018; Greenberg et al, 2016; Leong et al, 2019; Thakur et al, 2019; Consolati et al, 2020; Haren et al, 2020; Liu et al, 2020; Wieczorek et al, 2020b; Zimmermann et al, 2020; Zupa et al, 2020; Würtz et al, 2021). To address this hypothesis, we designed an NSEM assay to quantify the number of ring-shaped particles formed by γ-TuSC:Mzt1 and MGM. First, we identified 0.05% (w/v) glutaraldehyde as a suitable crosslinking concentration for MGM and γ-TuSC:Mzt1 (Suppl. Fig. 2). To assess whether we could quantitatively recover γ-TuSC:Mzt1 onto NSEM grids, we crosslinked γ-TuSC:Mzt1 (1 mg/ml concentration; approximately 400 nM). After quenching the reaction, we diluted the crosslinked samples such that the amount of γ-TuSC present in 100 µl ranged from 1×10^-12^ to 5×10^-12^ moles. We then centrifuged each dilution onto grids and visualised γ-TuSC:Mzt1 by NSEM (Suppl. Fig. 3A, B). Quantification from micrographs showed that the number of rings and ring fragments (i.e. with an arc of ≥ 180°, as estimated by eye) scaled linearly with moles of γ-TuSC:Mzt1 (R^2^ = 0.86; Fig. 1D). Notably, when compared to (non-centrifuged) samples with similar amounts of γ-TuSC:Mzt1 spotted directly onto grids, numbers of rings in centrifuged samples were increased ∼10-fold.

We then used our quantitative NSEM assay to determine whether, at lower concentrations, MGM could form more rings than γ-TuSC:Mzt1. We prepared both complexes at two different concentrations (20 nM and 400 nM), crosslinked them and compared the number of rings when equal numbers of moles were centrifuged onto grids. After imaging, particles were picked automatically, to reduce observer bias, and subjected to 2D classification (Fig. 1E). In four independent experiments, we observed no significant difference in the number of ring or ring fragments formed by the MGM and γ-TuSC:Mzt1 complexes, at either concentration (Fig. 1F). This suggests that the different microtubule-nucleation activities of MGM and γ-TuSC:Mzt1 in vitro (Leong et al, 2019) could not be explained by differences in ring formation.

### A minimal, microtubule-nucleating *S. pombe* γ-tubulin complex forms conical assemblies

Ring- and crescent-shaped particles of S. pombe MGM complex were observed in a previous study but were not further characterised (Leong et al, 2019). Using single particle analysis, we sought to determine the range of oligomeric states present in MGM samples (Fig. 2A, B). Within the concentration range of 5 nM to 50 nM (∼0.02 mg/ml to 0.2 mg/ml), particles ranged from 10 nm to 35 nm in their longest axis, with larger particles corresponding to rings and crescents. Smaller 10 nm particles likely correspond to MGM heterononamers (i.e. one molecule each of GCP2, GCP3, and Mzt1, and two molecules each of γ-tubulin, Mto1[bonsai], and Mto2) or smaller sub-complexes. Mto1[bonsai] forms dimers, and Mto2 has been shown to form a variety of oligomers in solution (Thakur et al, 2019), which may also contribute to sample heterogeneity.

**Figure 2:**
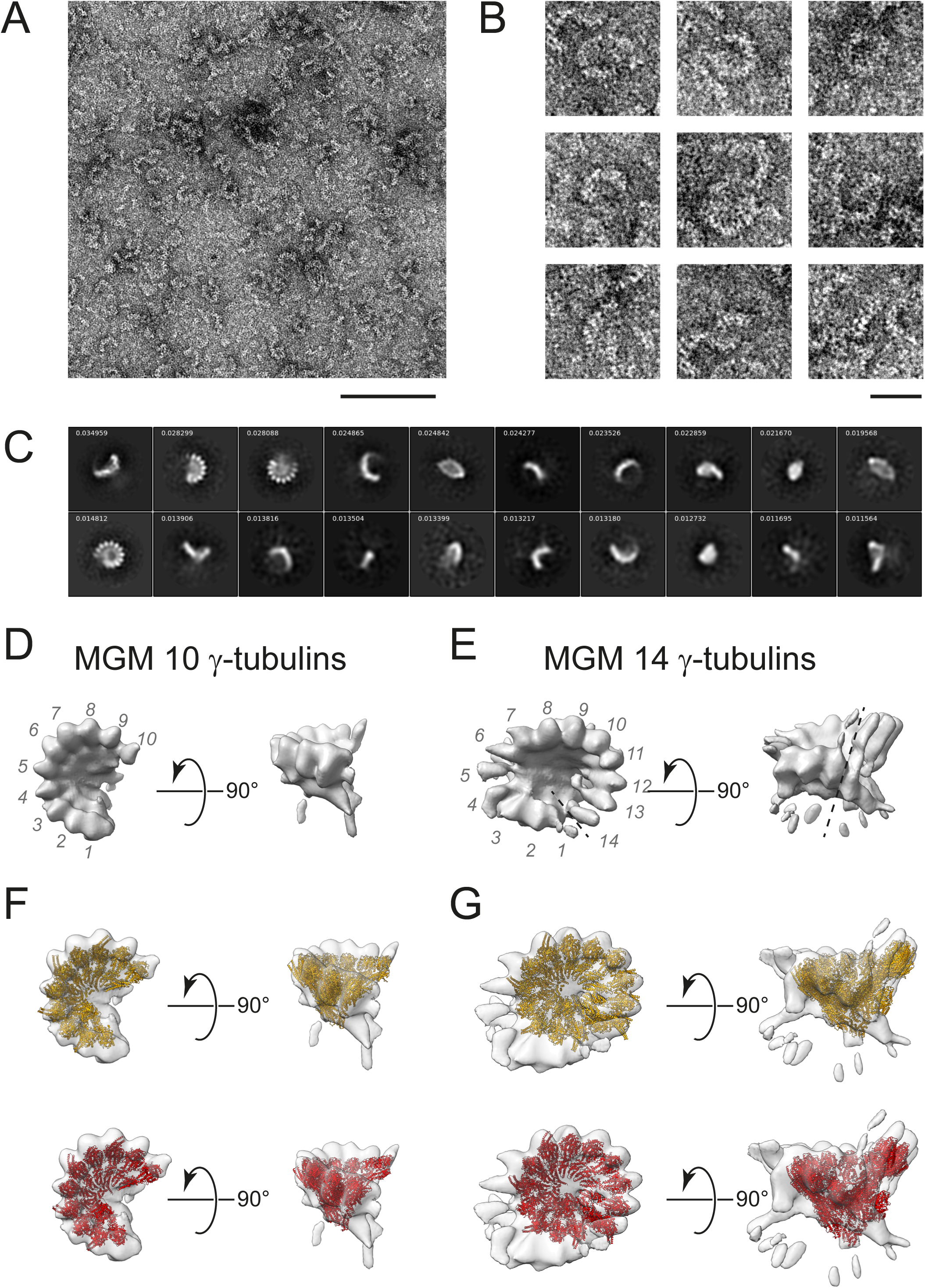
Negative stain EM reconstruction of MGM complex. **A)** Negative stain electron micrograph of MGM (50 nM concentration). **B)** Selected representative particles of MGM from multiple micrographs, as in A. Shapes observed: ring, spiral, crescent and rod. **C)** Top 20 most populated 2D class averages of crosslinked MGM from 2D classification of 15,206 particles using 200 classes and the EM algorithm. Numbers in the top left of each 2D class show the decimal fraction of particles in the class. D, E) 3D reconstructions of MGM showing the two identified oligomeric states, containing **(D)** 10 putative γ-tubulin molecules and **(E)** 14 putative γ-tubulin molecules. Numbers (gray italics) indicate γ-tubulin molecules. Dashed line in E marks the seam on the 14 γ-tubulin reconstruction. F, G) Comparisons of **(F)** 10 γ-tubulin and **(G)** 14 γ-tubulin (right) MGM reconstructions with models of native X. laevis open γ-TuRC (PDBID 6TF9; yellow) and H. sapiens closed γ-TuRC (PDBID 8VRK; red). Scale bars: (A) 150 nm; (B) 30 nm.

To limit MGM complex dissociation, we crosslinked samples with glutaraldehyde (Suppl. Fig. 2B, 2D) and carried out single particle analysis on NSEM images (Suppl. Fig. 4-6). After 2D classification, we separated classes of rings and crescents from classes of smaller particles (Suppl. Fig. 5) and reclassified in 2D. We observed well-defined densities on the circumference of both rings and crescents (Fig. 2C), leading us to interpret crescents as partial rings. Based on numerous similar structures (Kollman et al, 2010, 2015; Greenberg et al, 2016; Consolati et al, 2020; Huang et al, 2020; Liu et al, 2020; Wieczorek et al, 2020a, 2020b, 2021; Zimmermann et al, 2020; Zupa et al, 2020; Brilot et al, 2021; Würtz et al, 2021, 2022; Aher et al, 2024; Brito et al, 2024; Dendooven et al, 2024; Serna et al, 2024, 2026; Xu et al, 2024), we interpret these as the positions of γ-tubulin. In some classes of rings, we also observed well-defined radial densities similar to the arrangement of GCPs 2-6 and the lumenal bridge that have been characterized previously in other γ-TuRCs (Consolati et al, 2020; Liu et al, 2020; Wieczorek et al, 2020b, 2021; Zimmermann et al, 2020; Würtz et al, 2021, 2022; Aher et al, 2024; Dendooven et al, 2024; Gao et al, 2024, 2025) (Fig. 1E and 2C, Suppl. Fig. 5A).

After 2D classification of MGM, 15,206 ring- and partial ring-shaped particles were used to generate 3D reconstructions, revealing two possible conical structures, each with a narrow apex and wide base, with different numbers of sub-complexes (Fig. 2D and E, Suppl. Fig. 6A, B). The predominant structure, containing 10 γ-tubulins around the base, was derived from 75.2% of particles (Fig. 2D, Suppl. Fig. 4, Suppl. Fig. 6A). From the same 15,206 particles it was also possible to reconstruct a species of MGM with 14 γ-tubulins, from 33.8% of the particles (Fig. 2E, Suppl. Fig. 4, Suppl. Fig. 6B).

Vertebrate γ-TuRC structures, which have 14 γ-tubulins, can be in a “closed” conformation, which matches the MT lattice and supports efficient MT nucleation, or in an “open” conformation, which does not (Liu et al, 2020; Wieczorek et al, 2020b; Zimmermann et al, 2020; Würtz et al, 2022; Aher et al, 2024; Brito et al, 2024; Dendooven et al, 2024). Because MGM is an efficient MT nucleator, we wanted to determine whether our MGM reconstruction more closely matched a closed or an open conformation. However, we found that closed and open vertebrate and yeast γ-TuRC structures showed similarly good fits to our 10 γ-tubulin and 14 γ-tubulin reconstructions (Fig. 2F, G; Suppl. Table 1); this likely reflects the low resolution of our reconstructions.

We also investigated the small rod-shaped particles present in some highly-populated 2D classes (Fig. 2C). Because these particles had a distinct density at one end, we hypothesized that these might represent individual γ-TuSC:Mzt1 complexes. However, attempts to generate a 3D reconstruction from these classes were unconvincing, as the reconstructions did not have similar dimensions to the 2D class averages.

### γ-TuSC assemblies without and with Mto1[bonsai] and Mto2 are structurally distinct

To gain insight into why γ-TuSC:Mzt1 samples are poor nucleators relative to MGM, we used NSEM to analyse γ-TuSC:Mzt1 particles cross-linked with glutaraldehyde (Suppl. Fig. 2A, C). Most 2D classes showed rings, although some classes showed pairs of rings (Fig. 3A; Suppl. Fig. 7, Suppl. Fig. 8A); the pairs of rings may be an artifact of crosslinking (Suppl. Fig. 2C). 2D classes containing pairs of rings were excluded from subsequent analyses. When compared with the MGM 2D classes, the centers of the γ-TuSC:Mzt1 particles were darkly stained (compare Fig. 2C and 3A), and very few 2D classes showed bright, punctate density around the edges of rings (compare Fig. 2C and 3A and Suppl. Fig. 5A and 8A). In addition, unlike MGM, many 2D classes of γ-TuSC:Mzt1 deviated from circular shapes, showing pretzel-like or rectangular shapes with apparent two-fold symmetry (Fig. 3A, Suppl. Fig. 8B).

**Figure 3:**
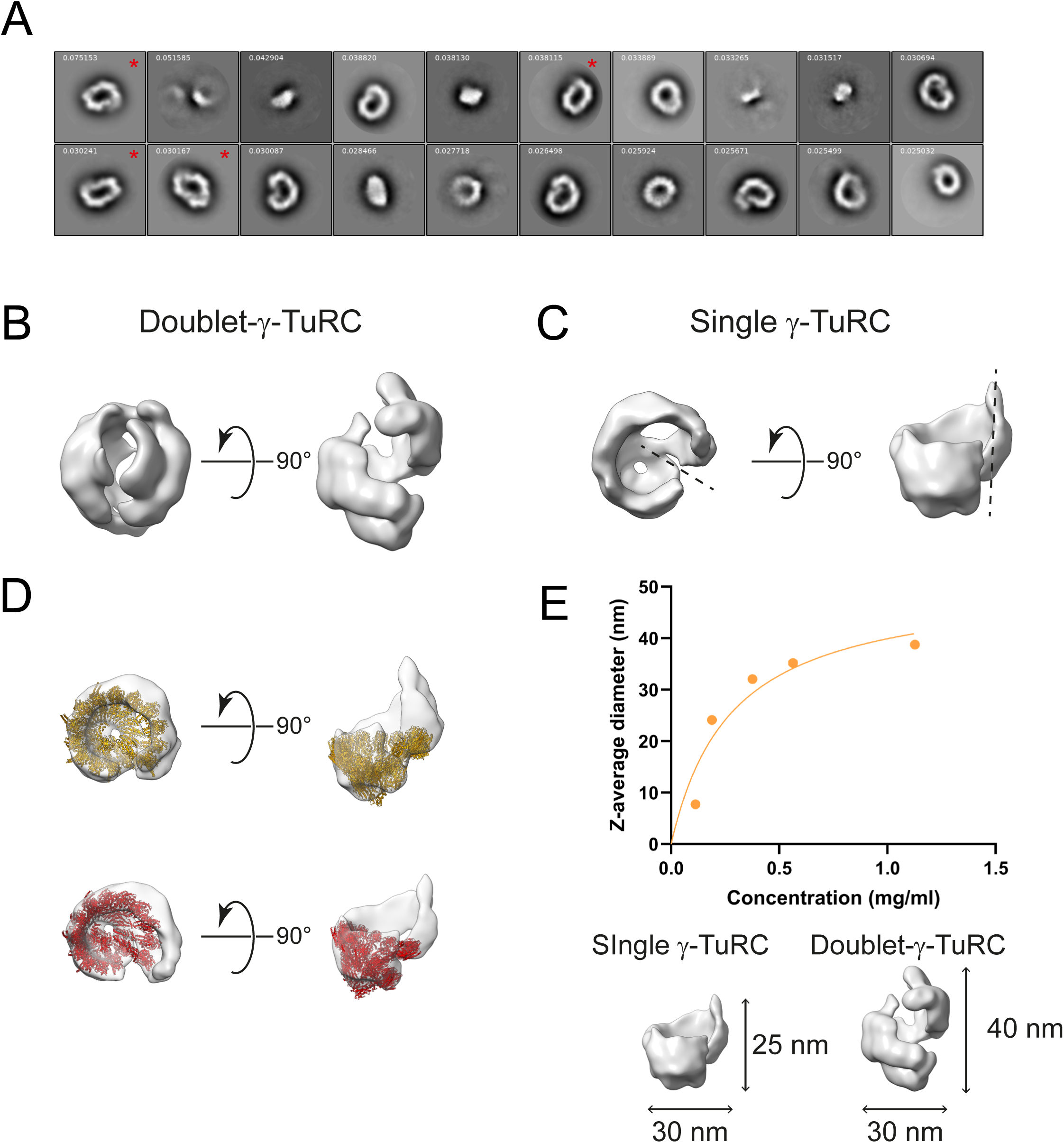
Negative stain EM reconstruction of γ-TuSC:Mzt1. **A)** Top 20 most populated 2D class averages of γ-TuSC:Mzt1 from 2D classification using 50 classes and the EM algorithm. Numbers in the top left of each 2D class show the decimal fraction of particles in the class. Classes containing rectangular shapes with apparent two-fold symmetry are marked with a red asterisk. **B, C)** 3D reconstructions of γ-TuSC:Mzt1 showing two identified conformations: a (B) doublet-γ-TuRC and a (C) single γ-TuRC. Dashed line in C marks the seam on the single γ-TuRC reconstruction. **D)** Comparison of single γ-TuRC γ-TuSC:Mzt1 reconstruction with models of native X. laevis open γ-TuRC (PDBID 6TF9; yellow) and H. sapiens closed γ-TuRC (PDBID 8VRK; red). **E)** Dynamic light scattering of γ-TuSC:Mzt1. Error bars representing standard deviation (calculated from three separate measurements of the same sample) are present but cannot be seen because they are smaller than the symbols. Molecular dimensions of the two reconstructions are shown below the graph.

To generate 3D reconstructions of γ-TuSC:Mzt1, we took two approaches (Suppl. Fig. 7). In one approach, we excluded only classes showing pairs of rings; in a second approach, we also excluded classes showing clear two-fold rotational symmetry (Suppl. Fig. 8B). When classes showing two-fold symmetry were retained, we observed a doublet γ-TuRC structure, in which two conical structures interact at their wide bases (which are interpreted to represent γ-tubulin; see above) (Fig. 3B, Suppl. Fig. 6C). However, when classes showing two-fold symmetry were excluded, we observed a single γ-TuRC ring (Fig. 3C, Suppl. Fig. 6D). Fitting known γ-TuRCs to the “single γ-TuRC” γ-TuSC:Mzt1 reconstruction gave good cross-correlation values (Suppl. Table 1), indicating that the global conical shape of this reconstruction broadly fits the geometry of a γ-TuRC (Fig. 3D). To investigate particle size in solution, we used dynamic light scattering (DLS) (Fig. 3E). We observed a monotonic relationship between hydrodynamic diameter (intensity-weighted Z-average) and γ-TuSC:Mzt1 concentration, with Z-average diameter reaching a plateau of 38.8 nm ± 0.1 nm (SEM) at the highest concentrations tested. This is consistent with doublet γ-TuRCs being present in the γ-TuSC:Mzt1 population.

The 3D reconstructions of MGM and γ-TuSC:Mzt1 suggested two non-exclusive explanations for the inability of γ-TuSC:Mzt1 to nucleate microtubules in vitro. First, if a significant proportion of γ-TuSC:Mzt1 complexes form doublets, then the γ-tubulins at the base of each conical structure become inaccessible for microtubule nucleation. Alternatively, we noticed that 3D reconstructions of single γ-TuRC γ-TuSC:Mzt1 complexes were poorly defined in comparison to MGM, with fewer distinct features, such as clearly defined γ-tubulin positions (Fig. 2D, E; Fig. 3C). This suggested that γ-TuSC:Mzt1 may be more structurally heterogeneous than MGM, such that γ-tubulins are not positioned correctly to template microtubule assembly. It is unlikely that differences between γ-TuSC:Mzt1 and MGM are due to dataset quality, because the γ-TuSC:Mzt1 and MGM reconstructions have a similar final resolution (19 Å for single γ-TuRC γ-TuSC:Mzt1, 18 Å for doublet γ-TuRC γ-TuSC:Mzt1, 28 Å for 10 γ-tubulin MGM, and 30 Å for 14 γ-tubulin MGM at Fourier Shell Correlation = 0.5), and more particles were used in reconstruction of γ-TuSC:Mzt1 compared with MGM (30,075 for single γ-TuRC γ-TuSC:Mzt1, 33,230 for doublet-γ-TuRC γ-TuSC:Mzt1, and 15,206 for MGM; Suppl. Figs. 4 and 7). Furthermore, the γ-TuSC:Mzt1 dataset and the two MGM datasets have similar data collection quality metrics (astigmatism mode 1000 Å for γ-TuSC:Mzt1 dataset and ∼800 Å & 900 Å for MGM datasets, contrast transfer function fit 5.5 Å and 7.0 Å & 5.5 Å, figure of merit 0.14 and 0.20 & 0.20, respectively).

To further explore structural differences between γ-TuSC:Mzt1 and MGM, we considered that 2D classifications from each complex may have features that would otherwise be lost in low-resolution 3D reconstructions. We therefore asked, if particle images from both complexes were combined prior to 2D classification, would they segregate into distinct classes after classification? To define appropriate images to combine, data from each sample (using the same datasets as were used previously) were processed to generate 100 2D classes, using the EM algorithm (Suppl. Fig. 9A, B). A ring class of γ-TuSC:Mzt1 was selected for separate comparisons against MGM rings, partial rings or side views (Suppl. Fig. 9B, C). In each comparison, all particles from the classes to be compared were combined in a new round of 2D classification that placed particles into only two classes (Class 1 and Class 2). We then counted the number of particles in each class originating from MGM vs γ-TuSC:Mzt1 datasets (Suppl. Fig. 9C). In analyses involving rings and partial rings, all particles in Class 1 were from γ-TuSC:Mzt1, and nearly all particles in Class 2 were from MGM, with only a few particles originating from γ-TuSC:Mzt1. As expected, γ-TuSC:Mzt1 rings did not co-classify with MGM side views. This suggests that γ-TuSC:Mzt1 and MGM rings are indeed distinct, even at the limited resolution of NSEM. Given the similar diameter and overall shape of the γ-TuSC:Mzt1 and MGM rings, it is likely that the MGM and γ-TuSC:Mzt1 segregate on the basis of features such as well-defined γ-tubulin densities and the presence vs absence of density at the center of the ring.

### Modelling local interactions within and between γ-TuSC and Mto1[bonsai]:Mto2 complexes

Given that our NSEM maps of MGM showed volumes that were unaccounted for after fitting of vertebrate cryo-EM γ-TuRC structures (Fig. 2F, G), and that MGM contains the additional proteins Mto1[bonsai] and Mto2, we next considered how these proteins associate with γ-TuSCs within MGM. We modelled interactions using AlphaFold3 and then compared models to crosslinking mass spectrometry (CLMS) data obtained from our previous characterisation of interactions within and between Mto1[bonsai] and Mto2 and γ-TuSCs (Leong et al, 2019; Thakur et al, 2019).

Focusing on local interactions, we first modelled a single γ-TuSC complex containing GCP2, GCP3, Mzt1 and two γ-tubulins (Suppl. Fig. 10; Suppl. Fig. 11 A-C). This complex closely resembled known γ-TuSC complexes (Lin et al, 2016a; Zupa et al, 2020; Brilot et al, 2021), although pTM and ipTM scores (0.59 and 0.55, respectively) were borderline for reliability. The pairwise aligned error (PAE) matrix is consistent with a well-folded structure and further suggested that the interaction between Mzt1 and the NTE of GCP3 (Leong et al, 2019) is independent of the rest of the complex (Suppl. Fig. 10; Suppl. Fig. 11C). We then compared the γ-TuSC AlphaFold3 model with our previous EDC crosslinking data (Leong et al, 2019). Taking into account an expected crosslink distance (Cα− Cα) of less than 12 Å (Merkley et al, 2014; Rivera-Santiago et al, 2015; Mintseris & Gygi, 2020) and known γ-TuSC flexibility (1.5 to 18.6 Å) (Zupa et al, 2020), we found that 73% of crosslinks (72 out of 99, within the structured region of γ-TuSC) fell within the expected distance (≤30 Å; Fig. 4A), showing that known interactions are appropriately recapitulated in the model.

**Figure 4:**
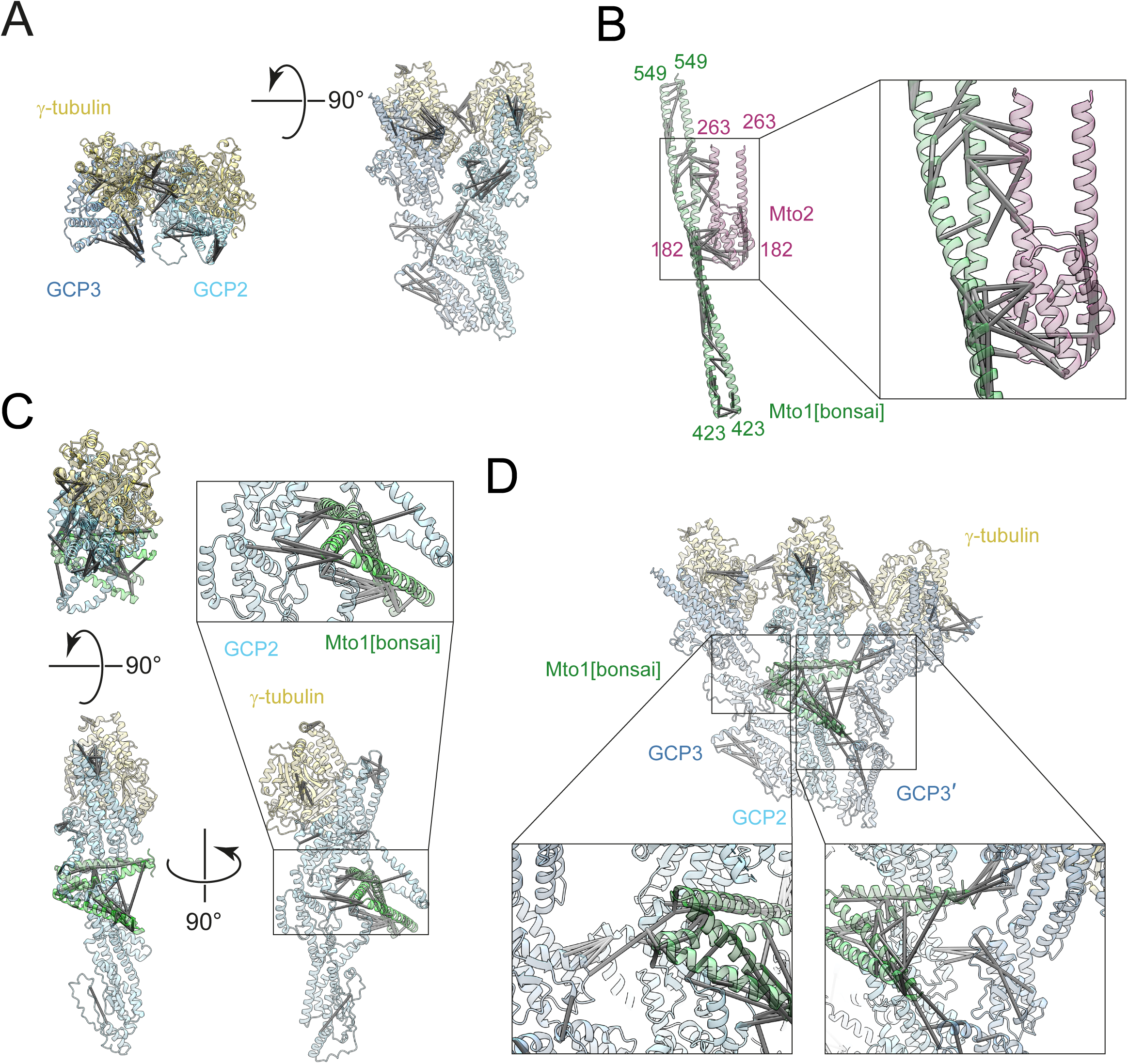
Mapping MGM crosslinks to MGM subcomplex AlphaFold3 models. Models for different MGM subcomplexes or fragments of subcomplexes, shown with experimentally determined chemical crosslinks (gray rods) having Cα - Cα distances <30 Å within the context of the models. **A)** γ-TuSC. **B)** Tetrameric fragment containing two copies of the Mto1[bonsai] coiled-coil region bound to an Mto2 twin cysteine domain (TCD) homodimer. Residue numbers shown are based on the residue numbering in full-length Mto1 and Mto2. **C)** Two molecules of the CM1 motif of Mto1[bonsai] (corresponding to residues 249-321 in full-length Mto1) bound to GCP2 plus one γ-tubulin. **D)** A “one-and-a-half” γ-TuSC multimer containing three γ-tubulins, two GCP3, one GCP2, and two Mto1 CM1 motifs. The additional GCP3 is labelled as GCP3’. The different shading of gray rods is a consequence of the angles of illumination used for displaying the models.

We next modelled an Mto1[bonsai]:Mto2 complex using a 2:2 stoichiometry (Suppl. Fig. 11D-F). Models generated by AlphaFold3 showed a high degree of variation (Suppl. Fig. 12A), perhaps because of long regions of coiled coil and few long-range interactions. Despite this variation, all models shared two common features: 1) homodimerization of Mto2 twin cysteine domains (TCD), which we previously determined experimentally (Thakur et al, 2019); and 2) interaction of helices 1 and 3 of the Mto2 TCD (residues 182-195 and 222-250, respectively) with a predicted coiled-coil segment of Mto1 (residues 476-504; Suppl. Fig. 11F, dashed boxes). These local interactions are well supported by CLMS data (Fig. 4B), and by prior experiments that mapped the region of Mto1 required for Mto2 binding in vitro (residues 476-549; (Samejima et al, 2005)).

Finally, we generated a model combining a single γ-TuSC with the Mto1[bonsai]:Mto2 complex (Suppl. Fig. 11G-I). Global features of the PAE matrix relating to the γ-TuSC:Mzt1 and Mto1[bonsai]:Mto2 complexes remained unchanged relative to the individual complexes (Supp. Fig. 11 C, F and I, Supp. Fig. 12B). In addition, a new interaction appeared, corresponding to a portion of the Mto1 interacting with GCP2 (Supp. Fig. 11I, dashed boxes). Specifically, the AlphaFold3 prediction placed the CM1 regions of the two Mto1[bonsai] molecules as a dimer at the hinge region of GCP2, with the first half of the CM1 regions most closely associated with GCP2 (Fig. 4C and Suppl. Fig. 11 G-I). This correlates well with CLMS data (Fig. 4C) and with prior analysis of mutations within the CM1 region in vivo (Samejima et al, 2008). It is also consistent with positions of CM1 regions in γ-TuSC/γ-TuRC proteins in S. scrofa and S. cerevisiae (Kollman et al, 2015; Greenberg et al, 2016; Brilot et al, 2021; Serna et al, 2024, 2026; Xu et al, 2024).

### Modelling of MGM complexes suggests an outer mantle of Mto1:Mto2

Although many crosslinks satisfied crosslinker-associated distance constraints in individual models of γ-TuSC:Mzt1 and Mto1[bonsai]:Mto2, other crosslinks violated these constraints (Suppl. Fig. 13). However, because γ-TuRCs comprise multiple copies of γ-TuSCs, we recognized that some crosslinks should be interpreted in the context of a multimeric structure. We therefore generated a “one-and-a-half” γ-TuSC model, containing three γ-tubulins, two GCP3s, one GCP2, and two Mto1 CM1 motifs, based on the GCP geometry within the native spindle pole body S. cerevisiae microtubule bound γ-TuRC (Dendooven et al, 2024). In this model, many of the previously violating crosslinks were now positioned at the inter-γ-TuSC interface, within the expected crosslink distance (Fig. 4D).

To assess how parts of Mto1[bonsai] and Mto2 other than the Mto1 CM1 region may associate with the core γ-TuRC ring, we next wanted to map the position of crosslinks from GCP2 and/or GCP3 to Mto1[bonsai] and/or Mto2 in the context of a γ-TuRC model. We constructed a 10 γ-tubulin γ-TuRC model by fitting the native microtubule-bound S. cerevisiae γ-TuRC (PDB ID – 8QV2) into our 10 γ-tubulin MGM reconstruction. We then aligned five copies of our S. pombe AlphaFold3 γ-TuSC model (with GCP3-NTE:Mzt1 removed) with the S. cerevisiae model. To retain conserved lateral interaction interfaces between GRIP1 motifs of GCP2 and GCP3, we used GCP2 as a reference point for alignment. When residues of γ-TuSC proteins that crosslink to Mto1[bonsai] and Mto2 were highlighted in the 10 γ-tubulin γ-TuRC model, most of these residues mapped to the outer surface of the cone or to the interface between adjacent γ-TuSCs (Fig. 5A). This indicates not only that the CM1 motif of Mto1 is in close interaction with GCP2, but also that other portions of Mto1[bonsai] and Mto2 can interact over an extensive surface of the outer γ-TuSC ring.

**Figure 5:**
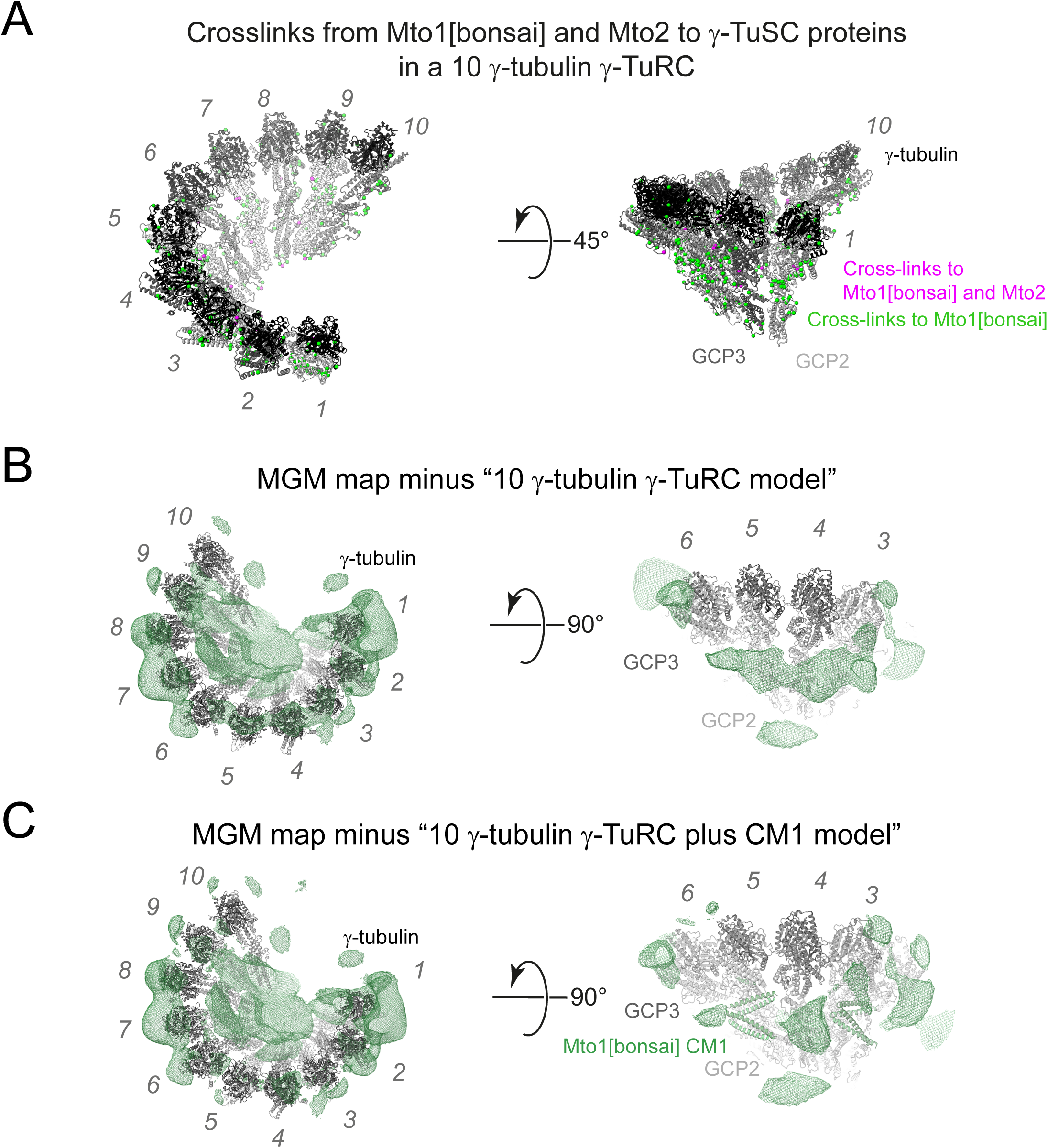
Integrative model building suggests that in MGM, Mto1[bonsai] and Mto2 form an outer mantle. **A)** Model of a 10 γ-tubulin γ-TuRC, with colored spheres indicating residues of γ-TuSC proteins that crosslink to Mto1[bonsai] (green) or to both Mto1[bonsai] and Mto2 (magenta). See text for details of modelling. **B)** Difference map (green wireframe) generated by subtracting the volume of the 10 γ-tubulin γ-TuRC model (black) from the MGM NSEM map. **C)** Difference map as in B, but subtracting the volume of the 10 γ-tubulin γ-TuRC model plus the Mto1 CM1 regions (Mto1 residues 249 -321) from the MGM NSEM map. CM1 is shown as green ribbon diagram. Numbers (gray italics) indicate γ-tubulin molecules.

To gain further insight into the arrangement of Mto1[bonsai]:Mto2 in MGM complexes, we subtracted the volume of our 10 γ-tubulin γ-TuRC model from the experimental MGM NSEM map (Fig. 5B). This revealed a ring of difference density, i.e. indicating missing mass, around the outside of the core γ-TuRC ring, broadly corresponding to the position of crosslinking sites described above (Fig. 5A). Reasoning that some of this extra density might correspond to the CM1 motif of Mto1, we added CM1 motifs to our model and repeated the subtraction of the model volume from the experimental density (Fig. 5C). The inclusion of CM1 motifs accounted for some of the missing mass, but difference density volumes remained, which likely represent mass contributions from other parts of Mto1[bonsai]:Mto2. While some of these volumes may be due to the Mto2 TCD homodimer domain, we do not have sufficient crosslinking data to determine how this domain associates with the core γ-TuRC ring, and thus the TCD homodimer domain, and its interaction with the dimeric coiled coil of Mto1 (Fig. 4B), could not be positioned unambiguously within the MGM model. Nevertheless, the remaining unaccounted for difference density suggest that the Mto1[bonsai]:Mto2 complexes contribute to a ring of density (a “mantle”) around the outside of the core γ-TuRC ring.

## Discussion

What makes MGM a good microtubule nucleator in vitro (Leong et al, 2019), while γ-TuSC:Mzt1 is a poor nucleator? Here we have addressed this question by asking what types of structure(s) can be made by MGM, and how this is changed in γ-TuSC:Mzt1, when Mto1[bonsai] and Mto2 are absent. We find that both MGM and γ-TuSC:Mzt1 can form rings, with similar efficiency, indicating that ring formation alone is not sufficient for microtubule nucleation. However, we also find that in MGM, positions of γ-tubulins align well across particles, while this is not the case in γ-TuSC:Mzt1. This implies that MGM has a more regular geometry than γ-TuSC:Mzt1. The tighter positioning of the γ-tubulins may be sufficient to template MTs. Given that the only difference between these complexes is the presence or absence of the Mto1[bonsai]:Mto2 complex, it seems likely that the Mto proteins rigidify the γ-TuSCs, resulting in a more regular γ-tubulin geometry (see below).

The majority of MGM particles have only ten γ-tubulins, while a smaller proportion have fourteen, as expected for complete γ-TuRCs. We do not know whether either of these forms preferentially templates MT growth. However, experiments using X. laevis proteins have shown that four tubulin heterodimer interactions with γ-TuRC are sufficient to facilitate MT nucleation (Thawani et al, 2020). This indicates that partial assemblies may facilitate MT nucleation, and it is possible that both ten and fourteen γ-tubulin forms can contribute to MT growth. We also found that γ-TuSC:Mzt1 particles can form doublets (at least in vitro), in which two separate particles interact via the γ-tubulins at their prospective nucleating surfaces. This could sequester γ-tubulins and thus block template-based MT nucleation. However, because many γ-TuSC:Mzt1 particles are single rings, it seems likely that unsuitable ring geometry prevents γ-TuSC:Mzt1 from functioning as a MT nucleator.

MT nucleation by MGM fits with previous observations that single and multiple deletions of GCP4, GCP5 and GCP6 in S. pombe lead to qualitatively normal but quantitatively decreased MT nucleation in vivo (Anders et al, 2006; Anders & Sawin, 2011; Lynch et al, 2014). Recent structural characterization of H. sapiens and X. laevis γ-TuRCs has shown that γ-TuSC-like complexes of GCP4:GCP5 and GCP4:GCP6 associate in a salt-resistant core complex (Liu et al, 2020), leading to the idea that these proteins nucleate or facilitate γ-TuRC assembly. Our in vitro reconstitutions using only S. pombe GCP2:GCP3 γ-TuSCs, plus Mzt1 (with or without Mto1[bonsai] and Mto2), show that rings can form despite the absence of GCP4, GCP5 and GCP6, consistent with previous in vivo analyses (Venkatram et al, 2004; Anders et al, 2006; Anders & Sawin, 2011; Lynch et al, 2014). However, our assemblies are highly heterogeneous, suggesting that the additional non-essential GCPs may limit structural and compositional heterogeneity of γ-TuRCs.

How does the Mto1[bonsai]:Mto2 complex reorganize the geometry of γ-TuSCs within MGM relative to γ-TuSC:Mzt1? Using AlphaFold3 modelling and comparing with CLMS data, we were able to identify a high-confidence local interaction between the Mto1 CM1 motif and GCP2 that corresponds with prior genetic data (Samejima et al, 2008), in vitro binding data (Cota et al, 2016; Leong et al, 2019) and structures of mammalian and S. cerevisiae γ-TuRCs (Wieczorek et al, 2020a, 2020b; Brilot et al, 2021; Dendooven et al, 2024; Serna et al, 2024, 2026; Xu et al, 2024; Gao et al, 2025). Further comparison of models with CLMS data confirmed the position of a local interaction between Mto1[bonsai] and Mto2, consistent with previous interaction mapping by pulldown assay (Samejima et al, 2005). Importantly, integration of a 10 γ-tubulin MGM complex γ-TuRC model with CLMS data strongly suggests that Mto1[bonsai] and Mto2 are located on the outer surface of the MGM cone. Consistent with this, our MGM NSEM map includes volumes on this outer surface that are not accounted for in γ-TuRC models based exclusively on γ-TuSC:Mzt1 components. These volumes likely reflect parts of Mto1[bonsai] and Mto2 present in a mantle that contributes to MGM stability; however, we have not been able to define their orientation relative to the cone.

How might a mantle of Mto1[bonsai] and Mto2 contribute to MT nucleation capability? Several possibilities are suggested. First, insertion of the Mto1[bonsai] CM1 motif into the hinge region of GCP2 likely constrains the orientation of its GRIP1 and GRIP2 motifs. Second, lateral interactions of the CM1 motif with GCP3 could allow stabilisation of the orientations of its GRIP1 and GRIP2 motifs, further reducing structural variation. Third, Mto2 dimers may help stabilise Mto1[bonsai] coiled-coil segments. Finally, the outer mantle of Mto1[bonsai]:Mto2 could further constrain the position of GCP proteins. In relation to these last two points, it is noteworthy that, although not apparent in metazoans, Mto2 orthologs are found in many fungi (Borek et al, 2015), and thus the contribution of Mto1 and Mto2 to MGM stabilization is likely conserved. This may represent one of many possible mechanisms by which different types of eukaryotic cells organize γ-tubulin for microtubule nucleation.

## Methods

### Insect cell culture

Recombinant protein expression used the baculovirus-insect cell expression system, using Sf9 cells for baculovirus production and High Five cells (Life Technologies) for protein expression. Sf9 and High Five cells were maintained in Sf-900 II SFM and Express Five SFM media, respectively (Life Technologies). Adherent cultures were split 1:5 on reaching confluency. Suspension cultures were maintained between 1×10^6^ – 8×10^6^ cells/ml. High Five suspension cultures were supplemented with 25 µg/ml dextran sulphate (Sigma).

### Generation of bacmids for protein expression

Bacmid generation used plasmids previously described by Leong et al. (Leong et al, 2019). For expressing Mto1[bonsai]:Mto2, a bacmid was generated from pKS1225, which contains N-terminally 6xHis-tagged Mto1[bonsai] and untagged Mto2. For expressing γ-TuSC:Mzt1, bacmids were generated from pKS1794, pKS1788 and pKS1801, which contain two copies of γ-tubulin, plus one copy each of GCP2 and C-terminally MBP-6xHis-tagged GCP3 (pKS1794), GCP2 and γ-tubulin (pKS1788), and C-terminally 6xHis-tagged Mzt1 (pKS1801). Bacmids were constructed by integration into DH10 EMBacY E. coli, using blue-white selection to identify integrants, followed by bacmid purification (Leong et al, 2019).

### Baculovirus generation and protein expression

Baculovirus-based protein expression was performed exactly as described previously (Leong et al, 2019). Baculovirus stocks were generated and amplified using Sf9 cells, and proteins were expressed using High Five cells. For expression of Mto1[bonsai]:Mto2, V_2_ baculovirus stocks were used to infect High Five cells (2×10^6^ cell/ml) at a ratio of 1:80 virus stock: cell-culture medium. For expression of γ-TuSC:Mzt1, High Five cells were co-infected with equal volumes of V_2_ stocks derived from pKS1794, pKS1788 and pKS1801 (Leong et al, 2019), added sequentially at a ratio of 1:80 virus stock: cell-culture medium. This was done to ensure that untagged proteins were expressed in excess of MBP-6xHis-tagged GCP3. After 48 hours, cells were harvested by centrifugation, washed with PBS and snap-frozen in liquid nitrogen.

### Protein purification

All proteins were purified as previously described (Leong et al., 2019). Mto1 [bonsai]:Mto2 complex was purified using ammonium sulphate precipitation, nickel affinity chromatography and size exclusion chromatography. For γ-TuSC:Mzt1, affinity purification using amylose resin was used. For experiments involving glutaraldehyde crosslinking (see below), Mto1 [bonsai]:Mto2 was dialysed into H100 buffer (40 mM HEPES, 100 mM KCl, 1 mM EGTA, 1 mM MgCl_2_ pH 7.5).

### MGM Reconstitution

The MGM complex was reconstituted by mixing estimated equal molar ratios of Mto1[bonsai]:Mto2 and γ-TuSC:Mzt1 (assuming a ratio of 1:1:2:1:2:2 GCP2:GCP3: γ-tubulin:Mzt1:Mto1[bonsai]:Mto2 per MGM heterononamer) at 4°C for 2 hrs. Subsequently, MGM was purified by sucrose gradient ultracentrifugation (Beckman Coulter Optima™ MAX Ultracentrifuge) using a 5 %-20 % isokinetic sucrose gradient in H100 buffer at 259,000 RCF (55,000 rpm) in a swinging bucket rotor (TLS 55) for 90 mins at 4°C.

### Dynamic Light Scattering (DLS)

DLS measurements were carried out using a Malvern Zetasizer APS with wavelength 830 nm. Light scattering was monitored at a fixed angle of 90°. To calculate an accurate viscosity constant for each solution, the buffer components and their concentrations were accounted for. Data were derived from the correlation function of the scattered intensity to calculate a radius of gyration (Rg). The Rg was then used to estimate the size of the particles by modelling them as a solid sphere.

### Glutaraldehyde crosslinking

To determine optimal glutaraldehyde concentrations for crosslinking prior to NSEM, protein complexes were crosslinked using 0.01%-0.15% (w/v) glutaraldehyde and analyzed by SDS-PAGE and NSEM (Suppl. Fig. 2). Crosslinked proteins were separated by SDS-PAGE using 3-8 % Criterion XT Tris-Acetate precast protein gels (Bio-Rad, 3450131). Sample and grid preparation for NSEM were performed as described below. Based on this optimization, both MGM and γ-TuSC:Mzt1 protein complexes were crosslinked using 0.05 % (w/v) glutaraldehyde (5 mM) for 5 minutes on ice (crosslinking volume 1 µl). Crosslinking was quenched by addition of Tris-HCl pH 7.5 to a final concentration of 100 mM and incubation on ice for 1 hour. Subsequently, samples were diluted in H100 buffer as required.

### Centrifugation of protein onto electron microscopy grids

For the quantitative NSEM assay of MGM and γ-TuSC:Mzt1 protein complexes (Fig. 1D-F, Suppl. Fig. 3), NSEM assay grids were glow discharged for 60 seconds at 40 mA using a PELCO easiGlow glow discharge machine. The centrifuge tube apparatus was assembled as shown in Suppl. Fig. 3 with a Perspex cylinder placed inside a centrifuge tube. 50 µl of buffer was pipetted between the Perspex cylinder and the tube wall. A further 1 µl of buffer was pipetted onto the top of the Perspex cylinder, to ensure that the grids adhered to the surface, after which the grids were then added to the centrifuge tube. 100 µl of crosslinked and diluted protein was then layered over the grid in the assembled tube apparatus on ice (Suppl. Fig. 3). The solution was then centrifuged in a Beckman Coulter Optima™ MAX ultracentrifuge at 214,000 RCF (50,000 rpm) in a TLS-55 swinging bucket rotor at 4 °C for 15 mins. After centrifugation, 100 µl of protein solution was removed by pipetting. The Perspex cylinder was then removed and placed onto a KIMTECH tissue, and the grid was recovered. Excess protein was blotted away with Whatman 150mm filter paper (Cat No 1001-150). Grids were then stained as described below.

### Coating grids with extra carbon layer

To coat carbon-coated grids with an extra layer of carbon, grids were placed in a petri dish on filter paper with the carbon film facing upwards. The grids were then placed in the vacuum chamber of a sputter coater, and a sharpened carbon rod was inserted. The vacuum chamber was pumped for 90 minutes to a pressure of 4×10^-7^ Torr. To evaporate carbon onto the grids, a power setting of 4 was used and three evaporations of 2 seconds were carried out.

### Staining grids for negative stain electron microscopy (NSEM)

To prepare grids for NSEM, 3.05 mm copper grids (400 mesh) with carbon support film were glow discharged for 30 to 90 seconds using a PELCO easiGlow glow discharge machine. 4 µl of protein sample was placed on the carbon film and incubated at room temperature for 2 mins. Excess sample was blotted using Whatman 150mm filter paper (Cat No 1001-150). Grids were washed and blotted four times with 15 µl H100 buffer. Grids were stained with 15 µl 2% (w/v) uranyl acetate with immediate blotting and then incubated with a further 15 µl 2% (w/v) uranyl acetate for 1 min.

### Electron microscopy data collection

NSEM micrographs were acquired using either a JEOL TEM-1400 plus TEM electron microscope with OneView Camera or an FEI TECNAI F20 200 keV cryo-electron microscope with TVIPS F816 detector. For single particle analysis, images were acquired on the TECNAI F20, using SerialEM to automate data collection (two sessions for each sample). The following SerialEM settings were used: 200 kV, 2.56 Å per pixel, -1.5 µm target defocus. A grid square was mapped in the SerialEM software, and 412 to 1072 non-overlapping micrographs were recorded.

### Single particle analysis

For each dataset, micrographs were initially viewed using EMAN2 (Tang et al, 2007). Images that contained the copper grid bar and/or had visual artefacts were excluded from the dataset. Particle dimensions were measured in ImageJ (Schindelin et al, 2012; Schneider et al, 2012). MGM and γ-TuSC:Mzt1 rings were determined to have a maximal dimension of ∼35 nm (140 pixels); therefore, a particle size of 35 nm and box size of 352 pixels (90.1 nm) were used for particle picking. For MGM and γ-TuSC:Mzt1 datasets, the EMAN2 Gaussian particle picker was used with the following settings: Use Variance = ON, Threshold low = 1, Threshold high = 2, Gaussian width = 1. After particles were picked, a subset of micrographs was checked to assess particle picking accuracy. The boxes were centred using the “ACF center all” command. Boxfiles were exported using the Generate Output (e2boxer.py) interface using the settings: boxsize = 352 pixels, ptclesize = 35 nm, apix = 2.56, with “only write_dbbox” ticked.

Subsequent processing used Relion v4.0. Contrast transfer function (CTF) estimation was carried out using GCTF with the Ignore ‘Searches’ parameter set to No and the Perform equi-phase averaging parameter set to No. Power spectra of the micrographs were used to identify and remove images with signs of drift. Particles from the remaining micrographs were then extracted with a particle box size of 352 pixels (90.1 nm). After particle extraction, there were 44,998 particles for the γ-TuSC:Mzt1 dataset and 29,789 particles for the MGM dataset. During 2D classification, both the VDAM and EM algorithms were tested with the following settings consistent across all jobs: regularisation parameter T = 2, mask diameter = 720 Å, limit resolution E-step to = 15 Å, Center class averages = Yes. Initially, six 2D classifications using 50, 100 and 200 classes with both the EM algorithm (25 iterations) and VDAM (200 minibatches) were made. After 2D classification, subsets of particles were selected, picking only from classes that either had an arc of at least 180° or were considered to be side views of the complex. Subsequently, initial models were generated using the 3D initial model job with all settings set to default unless otherwise specified. For all 3D reconstructions, only those of good quality were taken forward, as indicated by pointed arrows in Suppl. Figs. 4 and 7. 3D reconstructions that did not meet the quality criteria (and were not taken forward) are indicated by flat headed arrows in the same figures. Quality of the reconstructions was determined by visually inspecting the reconstructions in ChimeraX. Criteria for good reconstructions included 1) not having any misfitting artefacts such as streaking or slicing, which would indicate misfitting of a plane in Fourier space; 2) having identifiable structural features such as γ-tubulin- or GCP-sized densities that were roughly the correct size; and 3) having a similar structure to known γ-TuRCs or sections of known γ-TuRCs. After this, 3D classification was carried out using a 3D initial model as a reference map and a subset selection of particles as an input STAR file. For 3D classification, “limit resolution E-step” to was set to 15 Å; all other settings were default. Favourable reconstructions generated during 3D classification were then used in a 3D auto-refine job. 3D auto-refine jobs were run with default settings.

### Comparison of γ-TuSC:Mzt1 and MGM ring particles from 2D classification

To compare γ-TuSC:Mzt1 and MGM datasets, Relion v4.0 was used. 2D classification was carried out with 100 classes using the EM algorithm (Suppl. Fig. 9A). After 2D classification, three separate pairwise comparisons were performed between γ-TuSC:Mzt1 particles from the most populated ring class and particles from three different classes of MGM: rings; partial rings; and side views (Suppl. Fig. 9B). Particles from the γ-TuSC:Mzt1 ring class were independently pooled with particles from the three MGM classes. MGM partial rings and side views were included as controls that would be expected to segregate cleanly from γ-TuSC:Mzt1 rings. 2D classification was then used to segregate the pooled particles into two classes (Suppl. Fig. 9C). Each class was saved individually using the Subset Selection job. The number of particles derived from MGM datasets and γ-TuSC:Mzt1 datasets were identified by counting input file names using grep commands in unix (e.g. grep -c G-TuSCMzt1 particles.star).

### Structural alignments of MGM maps and known structures

For all structural alignments of our NSEM-derived maps with existing γ-TuRC models, ChimeraX 1.8 (Goddard et al, 2018) was used. γ-TuSC:Mzt1 and MGM maps were dusted to remove any disconnected densities smaller than the corresponding map resolution (45Å for the 10 γ-tubulin MGM map, 42Å for the 14 γ-tubulin MGM map, 41Å for the γ-TuSC:Mzt1 map). The MGM and γ-TuSC:Mzt1 maps were compared with the native X. laevis γ-TuRC (PDB ID 6TF9) (Liu et al, 2020), recombinant H. sapiens γ-TuRC (PDB ID 7QJ5 and 7AS4) (Zimmermann et al, 2020; Würtz et al, 2022), native H. sapiens γ-TuRC (PDB ID 6V6S) (Wieczorek et al, 2020b), recombinant H. sapiens microtubule bound γ-TuRC (PDB ID 8VRK) (Aher et al, 2024), native spindle pole body S. cerevisiae microtubule bound γ-TuRC (PDB ID 8QV2) (Aher et al, 2024) and the recombinant H. sapiens microtubule bound fully closed γ-TuRC (Brito et al., 2024, see associated supplementary dataset S1). To align models, they were first manually moved to overlap with the MGM or γ-TuSC:Mzt1 map, subsequently the Fit in Map function was used to optimise the fit. To determine correlation values, a map was simulated from atoms in the PDB models to the same resolution as the γ-TuSC:Mzt1 or MGM map to which it was being compared (10 γ-tubulin MGM map = 45Å, 14 γ-tubulin MGM map = 42Å, γ-TuSC:Mzt1 map = 41Å). The following “fit in map” settings were selected: “Use only data above the contour level from first map”, “optimise for correlation”, “allow rotation and shift”, and “move whole molecules”.

### AlphaFold3 structure prediction

Structures were predicted using AlphaFold3 on the AlphaFold Server (Abramson et al, 2024). Sequences used for structural prediction were derived from plasmids used for the purification of MGM (Leong et al, 2019) for SpGCP2, SpGCP3, Mto1[bonsai], Mto2, Mzt1 and γ-tubulin; for full length Mto1, the sequence from Pombase (Harris et al, 2022) was used. To predict structures of multimers, the appropriate number of copies of the sequence were selected in the AlphaFold3 submission.

### AlphaFold3 complex modelling and crosslink mapping

MGM and γ-TuSC:Mzt1 model building was carried out in UCSF ChimeraX 1.8 (Goddard et al, 2018; Pettersen et al, 2021; Meng et al, 2023). Models of S. pombe proteins were generated using AlphaFold3 as above. During model building and volume subtractions, we considered structural information from previous characterisations of C. albicans and S. cerevisiae γ-TuSCs and γ-TuRCs, X. laevis, H. sapiens and S. scrofa γ-TuRCs as well as previous studies of S. pombe γ-TuRC proteins (Oegema et al, 1999; Gunawardane et al, 2000b; Choy et al, 2009; Kollman et al, 2010, 2011, 2015; Guillet et al, 2011; Lynch et al, 2014; Leong et al, 2019; Thakur et al, 2019; Consolati et al, 2020; Liu et al, 2020; Wieczorek et al, 2020a, 2020b, 2021; Zimmermann et al, 2020; Zupa et al, 2020, 2021; Würtz et al, 2021; Serna et al, 2024, 2026; Xu et al, 2024).

For visualisation in Fig. 4, flexible linker regions and model components not required for illustrating the relevant crosslinks were omitted from the AlphaFold3 models by deleting corresponding atoms. These modifications were performed after model prediction to aid interpretation of crosslinks mapped to models.

A GCP3-NTE:Mzt1 module was generated from the γ-TuSC:Mzt1 heteropentamer by deleting both γ-tubulins, GCP2, and GCP3 residues 109-832, while retaining the complete Mzt1 subunit (Suppl. Fig. 10).

For Fig. 4A, a γ-TuSC:Mzt1 heteropentamer containing GCP2, GCP3, γ-tubulin and Mzt1 in a 1:1:2:1 ratio was predicted using AlphaFold3 (Fig. 4A and Suppl. Fig. 11A). For visualization, GCP2 residues 1-71, GCP3 residues 1-194 and all Mzt1 residues were deleted.

For Fig. 4B, an Mto1[bonsai]:Mto2 heterotetramer using a 2:2 ratio was predicted using AlphaFold3. For visualization, Mto1 residues 1-422 and 550-1115 (using full-length numbering for Mto1), and Mto2 residues 1-181 and 264-397 were deleted.

For Fig. 4C, an MGM heterononamer containing GCP2, GCP3, γ-tubulin, Mzt1, Mto1[bonsai] and Mto2 in a 1:1:2:1:2:2 ratio, was predicted using AlphaFold3. For visualization, one copy of γ-tubulin, Mto1 residues 1-248 and 322-1115, and all of GCP3, Mto2, Mzt1, were deleted.

For Fig. 4D, a model of γ-TuSC containing a neighbouring GCP3 and comprising GCP2, GCP3, γ-tubulin and Mto1[bonsai] in a 1:2:3:2 ratio was predicted using AlphaFold3. For visualization, GCP2 residues 1-71 and Mto1 residues 1-248 and 322-1115 were deleted.

The AlphaFold3 models were evaluated against experimental CLMS data to assess their quality. Crosslinks were exported as ChimeraX pseudobonds using XiView (Combe et al, 2024) (Fig. 4). For Fig. 4 A, C and D, MGM crosslinks were used (Leong et al, 2019). For Fig. 4B, Mto1[bonsai]:Mto2 multimer crosslinks were used (Thakur et al, 2019).

### MGM model building and map subtraction

To build an MGM model, models of the native S. cerevisiae spindle pole body γ-TuRC bound to microtubules (Dendooven et al, 2024) (PDB ID 8QV2) and of recombinant H. sapiens γ-TuRC (Würtz et al, 2022) (PDB ID 7QJ5) were fit into the 10 and 14 γ-tubulin MGM reconstructions using the ChimeraX “fit in map” tool. Multiple copies of the MGM heterononamer AlphaFold3 model were then aligned to 8QV2 using the matchmaker command in ChimeraX to align the GCP2 chains. 8QV2 was used for alignment as it had the highest cross correlation value to the map (Suppl. Table. 1). For Fig. 5A, to visualize where crosslinks from Mto1[bonsai] and/or Mto2 to γ-TuSC proteins are positioned on the γ-TuRC, Mto1[bonsai], Mto2, Mzt1, GCP2 residues 1-71 and 508-556, and GCP3 residues 1-194 were deleted. For Fig. 5B, a γ-TuRC model lacking Mto1[bonsai] CM1 was subtracted from the MGM NSEM map. For Fig. 5C, a model including Mto1[bonsai] CM1 was subtracted from the NSEM map. Subtraction was performed by generating a map based on the model at 28 Å, the resolution at the FSC=0.5 cutoff, then subtracting the simulated map from the experimental data map of MGM with 10 γ-tubulins.

## Data Accessibility

3D volumes for electron microscopy have been deposited in EMDB, with accession numbers EMD-59427 for the doublet γ-tubulin ring complex, EMD-59428 for the MGM complex – partial ring, EMD-59429 for the γ-tubulin ring complex – γ-TuSC:Mzt1 and EMD-59430 for the MGM complex – complete ring. AlphaFold3 models and reconstructions are available at DOI: 10.5281/zenodo.23010657.

## Author contributions (Using CRediT roles)

JDLC – Formal Analysis, Investigation, Visualization, Writing Original Draft, Review and Editing

SLL – Investigation, Formal Analysis, Validation

EML – Investigation

HCT – Investigation

KES – Conceptualization, Supervision, Funding Acquisition, Writing Original Draft, Review and Editing

AGC – Conceptualization, Supervision, Funding Acquisition, Writing Original Draft, Review and Editing

## Acknowledgements

We are grateful to Martin Singleton and Maartin Tuijtel and the EM facility for support with EM data collection and analysis. Data/Grid screening was acquired/performed in the cryoEM facility in School of Biological Sciences at the University of Edinburgh. The cryoEM facility was set up with funding from Wellcome (087658/Z/08/Z) and SULSA. The authors would like to acknowledge Stephen Mitchell at the School of Biological Sciences electron microscopy unit for assistance with EM. We acknowledge the support of a Wellcome Multi-User Equipment Grant (WT104915MA). JDLC was supported by the University of Edinburgh. AGC was supported by a Wellcome Senior Fellowship (200898). SLL, HCT, and KES were supported by a Wellcome Trust Investigator Award (210659 to KES). EML was supported by a PhD studentship from the Darwin Trust of Edinburgh. This work was carried out in the Wellcome Centre for Cell Biology, supported by core funding from Wellcome (203149). This work utilised the Edinburgh Protein Production Facility (EPPF) and the Centre Optical Instrumentation Laboratory, funded by Wellcome Core Grants 092076 and 203149

## Abbreviations

CLMS: Crosslinking mass spectrometry
CM1: Centrosomin motif 1
Cryo-EM: Cryo-electron microscopy
DLS: Dynamic light scattering
γ-TuC: γ-tubulin complex
γ-TuSC: γ-tubulin small complex
γ-TuRC: γ-tubulin ring complex
GCP: γ-tubulin complex protein (e.g. GCP2, GCP3, GCP4, GCP5, GCP6)
GRIP motif: γ-tubulin ring protein motif
Gtb1: γ-tubulin
Mto1: Microtubule organizer 1
Mto2: Microtubule organizer 2
Mzt1: Mitotic-spindle organizing protein 1
NSEM: Negative stain electron microscopy
NTE: N-terminal extension
Spc110: Spindle pole body component 110

