## Supplemental Figures for "*S. pombe* Mto1[bonsai]:Mto2 stabilizes γ-tubulin positions within the γ-TuRC"

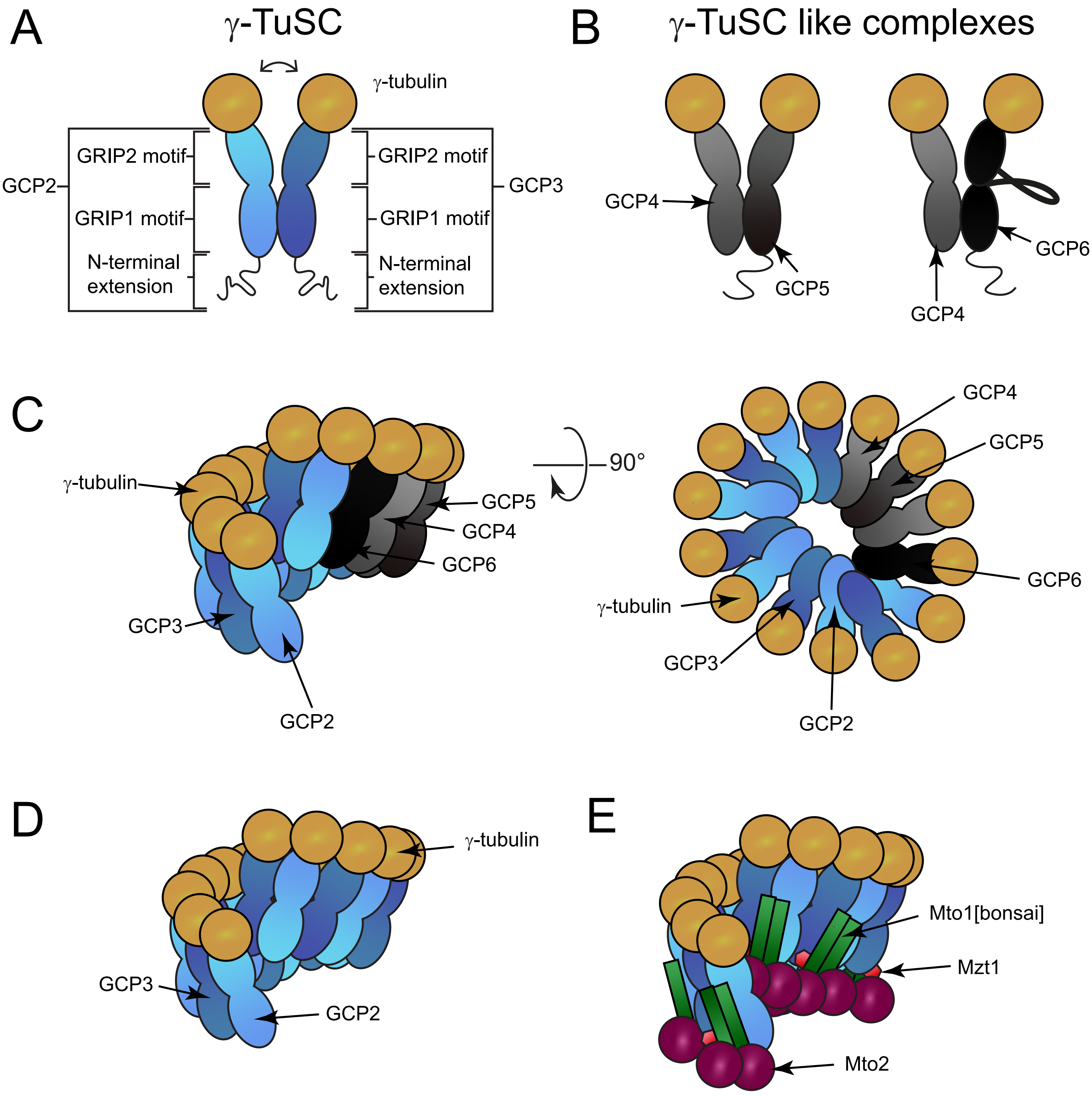


**Supplementary Figure 1: Schemas of γ-TuSCs, γ-TuSC-like complexes, and γ-TuRCs.**
**A**) γ-TuSC complexes. **B**) γ-TuSC-like complexes. **C**) Schema showing the positions and stoichiometries of metazoan γ-TuRC GCP proteins. **D**) Schema of the *S. cerevisiae* γ-TuRC. **E**) Schema of the *S. pombe* MGM complex.

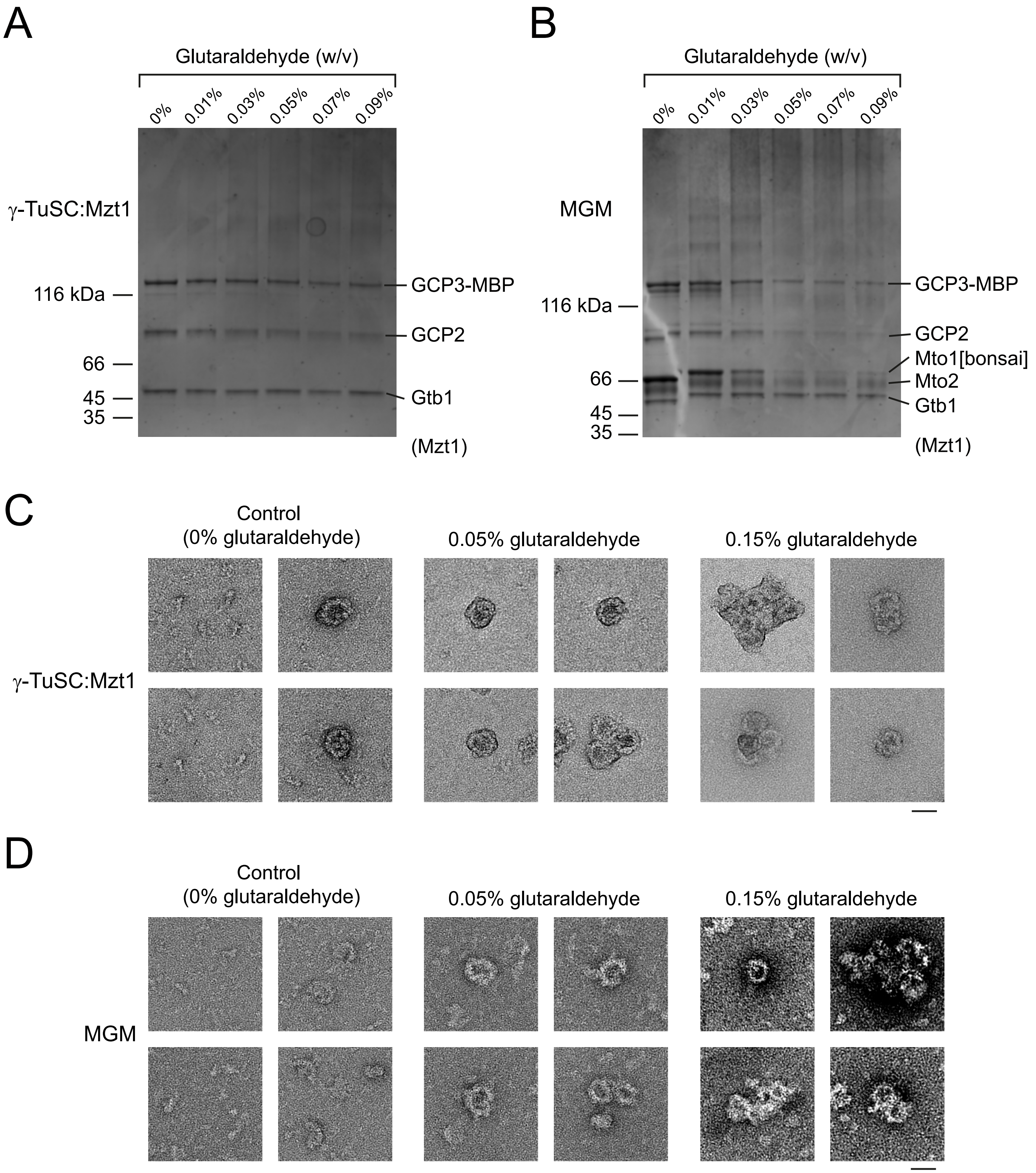


**Supplementary Figure 2: Glutaraldehyde crosslinking optimization of γ-TuSC:Mzt1 and MGM.**
**A, B**) Coomassie-stained SDS-PAGE of glutaraldehyde crosslinked **A**) γ-TuSC:Mzt1 and **B**) MGM. Mzt1 is present in the sample but is too small to be seen on the gel. **C, D)** Regions of negative stain electron micrographs showing different particle shapes identified in glutaraldehyde crosslinked **C**) γ-TuSC:Mzt1 and **D**) MGM. Scale bars, 30 nm.

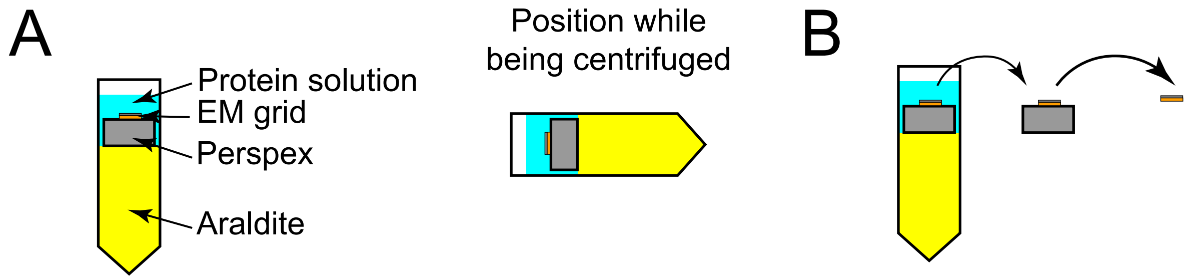


**Supplementary Figure 3: Quantitative negative stain electron microscopy assay tube design.**
**A**) Design of centrifuge tube used in a swinging bucket rotor to enrich protein on negative stain electron microscopy grids. **B**) Schematic of how grids are removed from centrifuge tubes after protein has been centrifuged onto the grids.

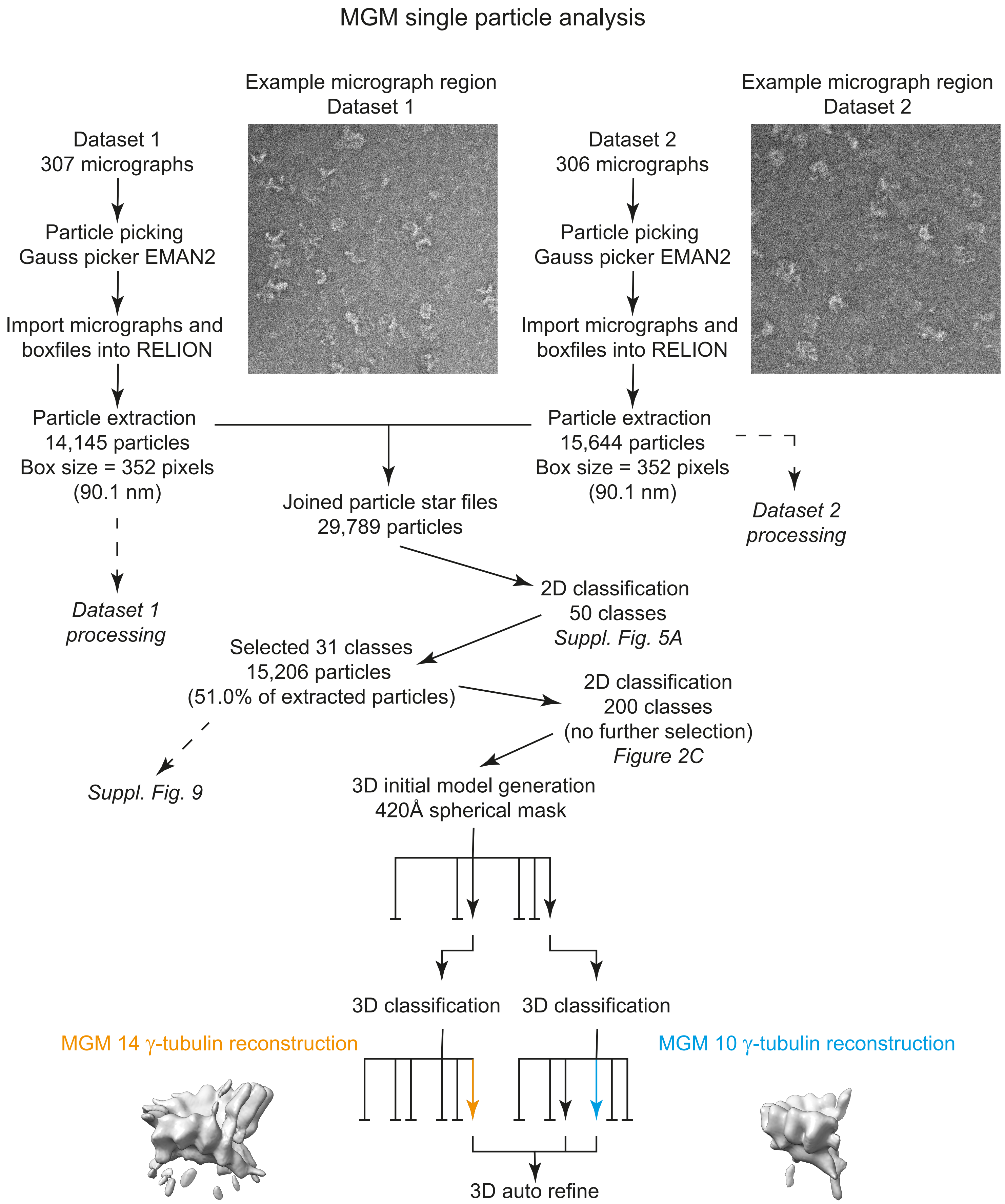


**Supplementary Figure 4: Data processing workflow for the single particle analysis of MGM.** Arrow-headed lines represent classes or particles which are used for further jobs, while bar-headed lines represent classes that were not taken forward. For 3D initial model jobs and 3D classification, job arrows are grouped into clusters, with each cluster representing a job. For each job, the leftmost class is class001 with classes ascending as they progress to the right through a cluster. 3D classes that were used to generate the 10 and 14 $\gamma$-tubulin MGM reconstructions in Figure 2D and E are shown in cyan and orange respectively. There are particles common to both reconstructions.

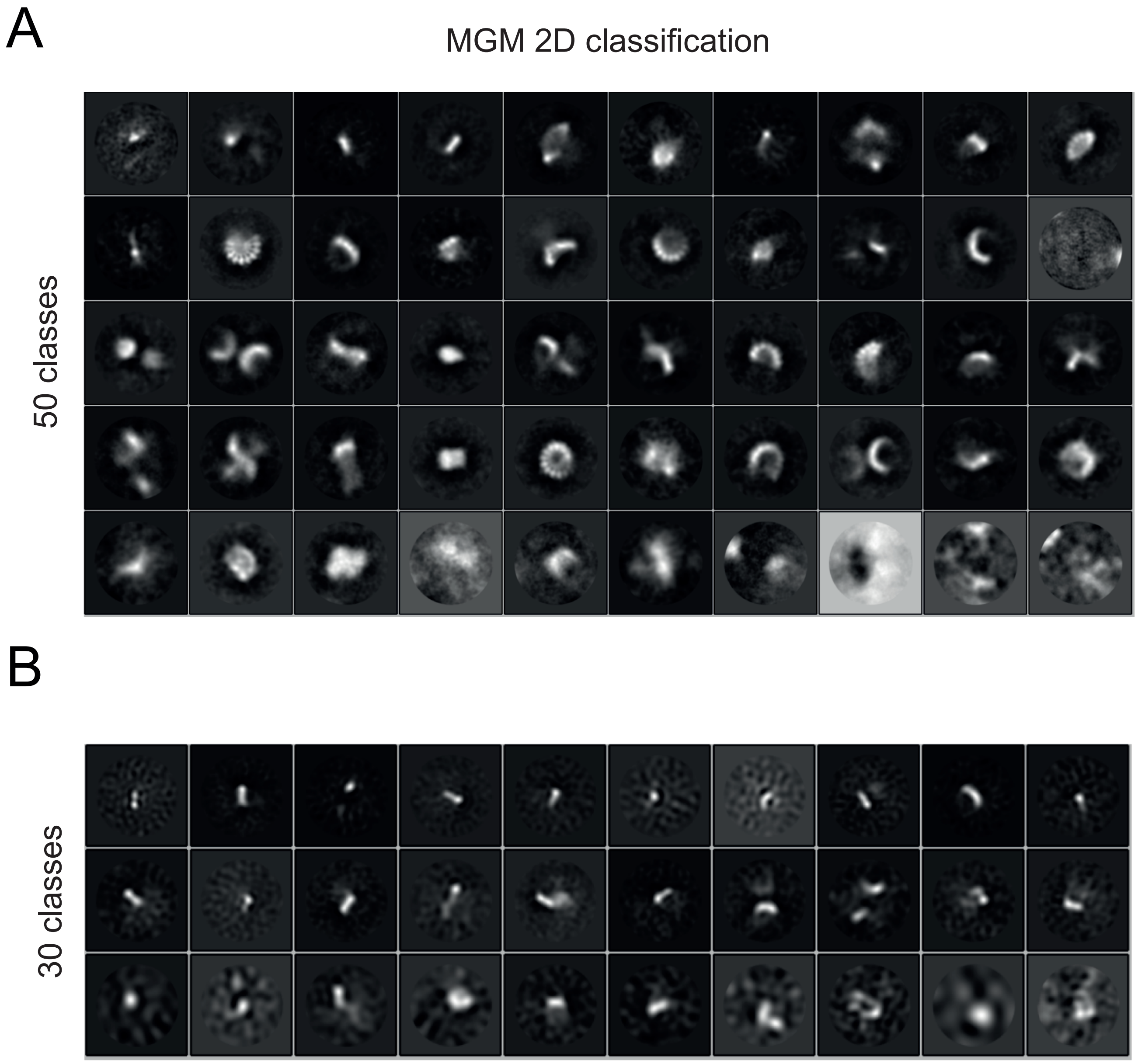


**Supplementary Figure 5: MGM 2D classification.**
**A**) 2D classification of MGM particles using the EM algorithm. **B**) Rod-shaped 2D classes from the MGM dataset. Classes are ordered by the number of particles contributing to each class, with the top-left class containing the most particles, and then row-by-row, with the bottom-right class containing the fewest particles.

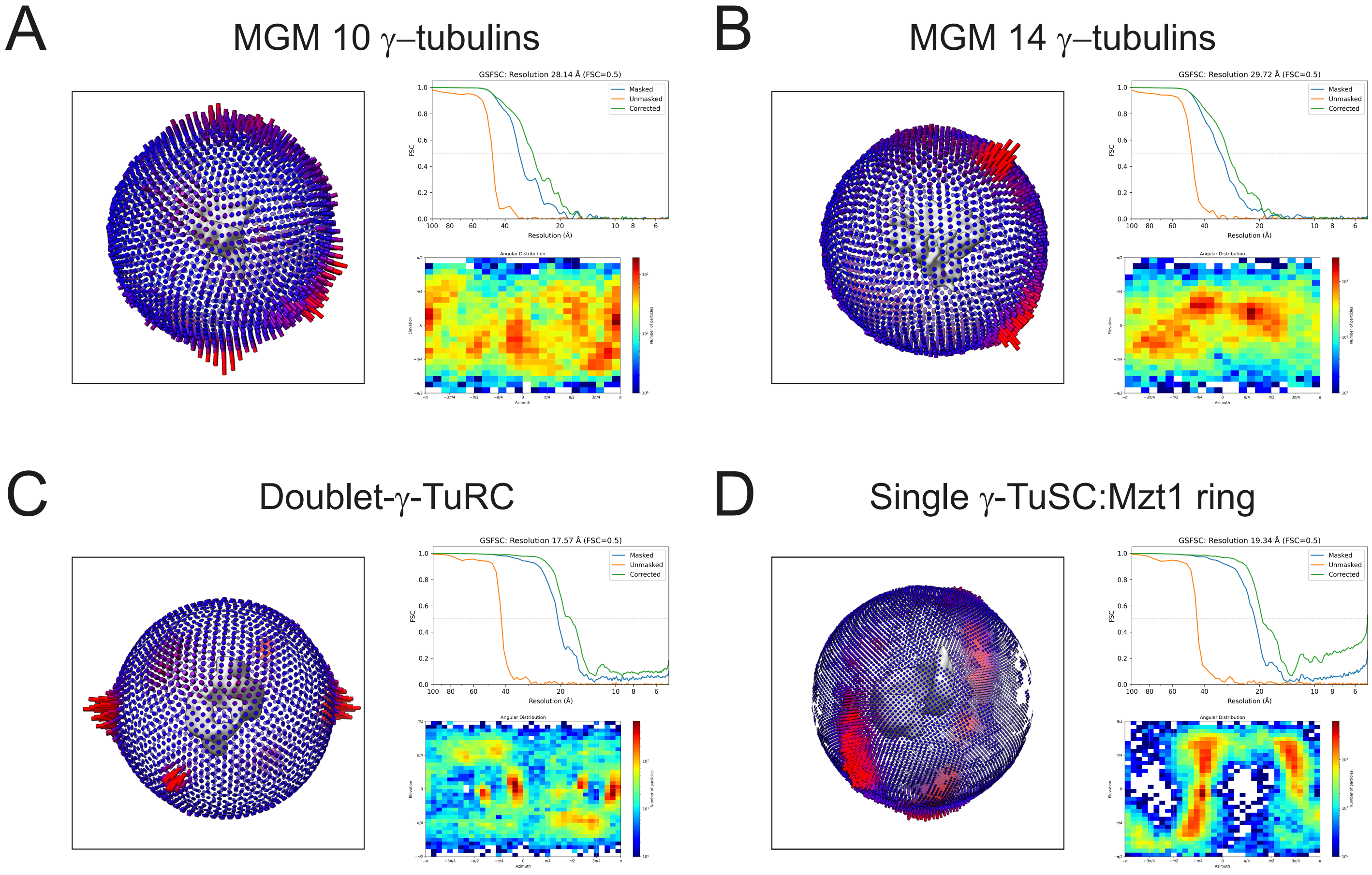


**Supplementary Figure 6: Reconstruction validation and angular distributions of particles.** Validation for 3D reconstructions of **A**) MGM containing 10 γ-tubulins, **B**) MGM containing 14 γ-tubulins, **C**) doublet γ-TuSC:Mzt1 ring and **D**) singlet γ-TuRC. Each panel shows a 3D angular distribution diagram on the left, with the NSEM map (gray) at the center, surrounded by a bar chart with the number of particles contributing to each angle in the reconstruction. The bar chart is colored based on the number of particles contributing to a given angle, ascending from blue to purple to red. The top right of each panel shows gold-standard Fourier Shell Correlation (GSFSC) plots for unmasked, masked and corrected curves. The bottom right of each panel shows angular distribution plotted as a 2D heat map, with red indicating a large number of particles and blue indicating a small number.

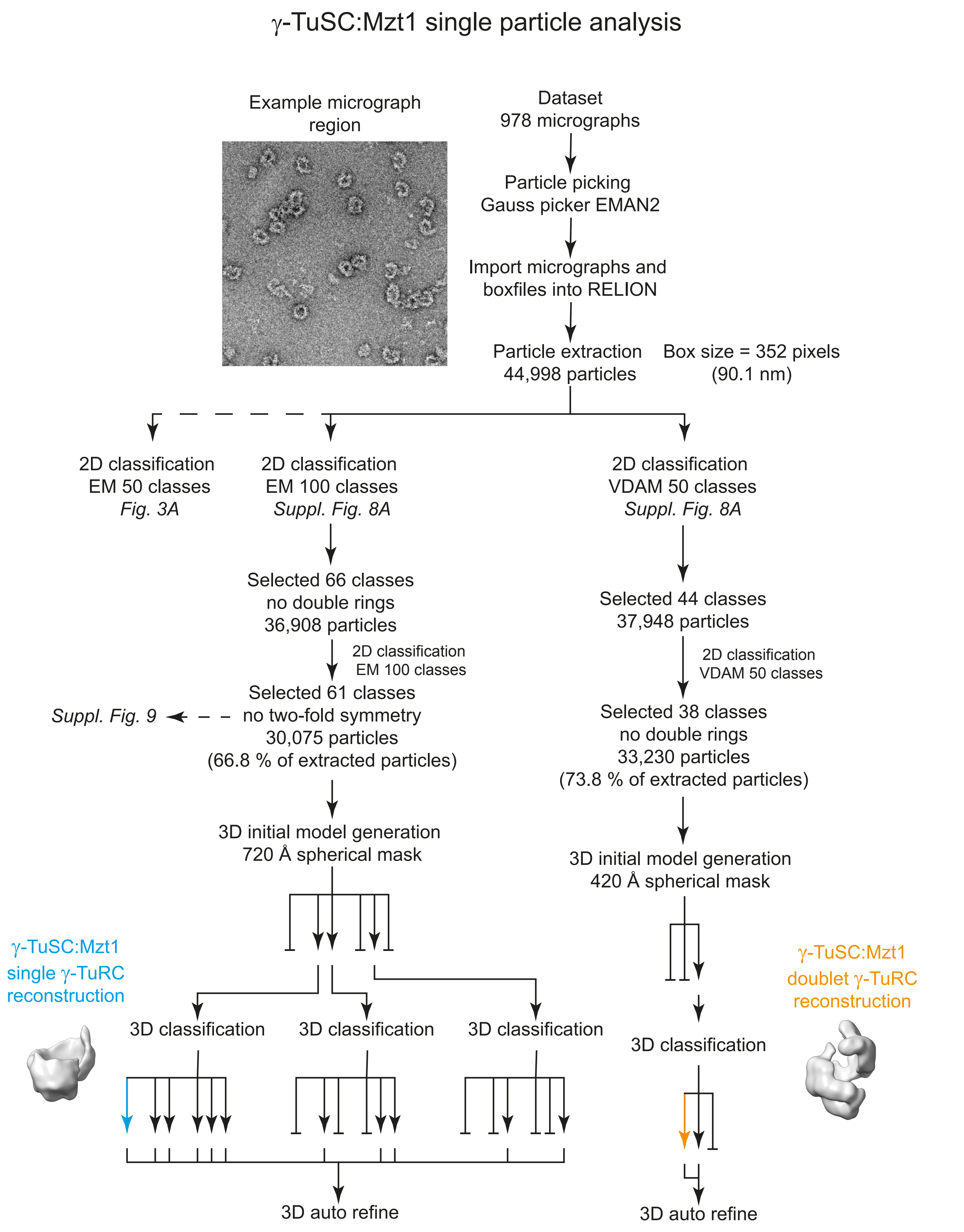


**Supplementary Figure 7: Data processing workflow for the single particle analysis of γ-TuSC:Mzt1.** Arrow-headed lines represent classes or particles that were used for further jobs, while bar-headed lines represent classes that were not taken forward. For 3D initial model jobs and 3D classification jobs, arrows are grouped into clusters with each cluster representing a job. For each job the leftmost class is class001 with classes ascending as they progress to the right through a cluster. 3D classes that were used to generate the single γ-TuRC and doublet-γ-TuRC reconstructions in Figure 3B are shown in cyan and orange.

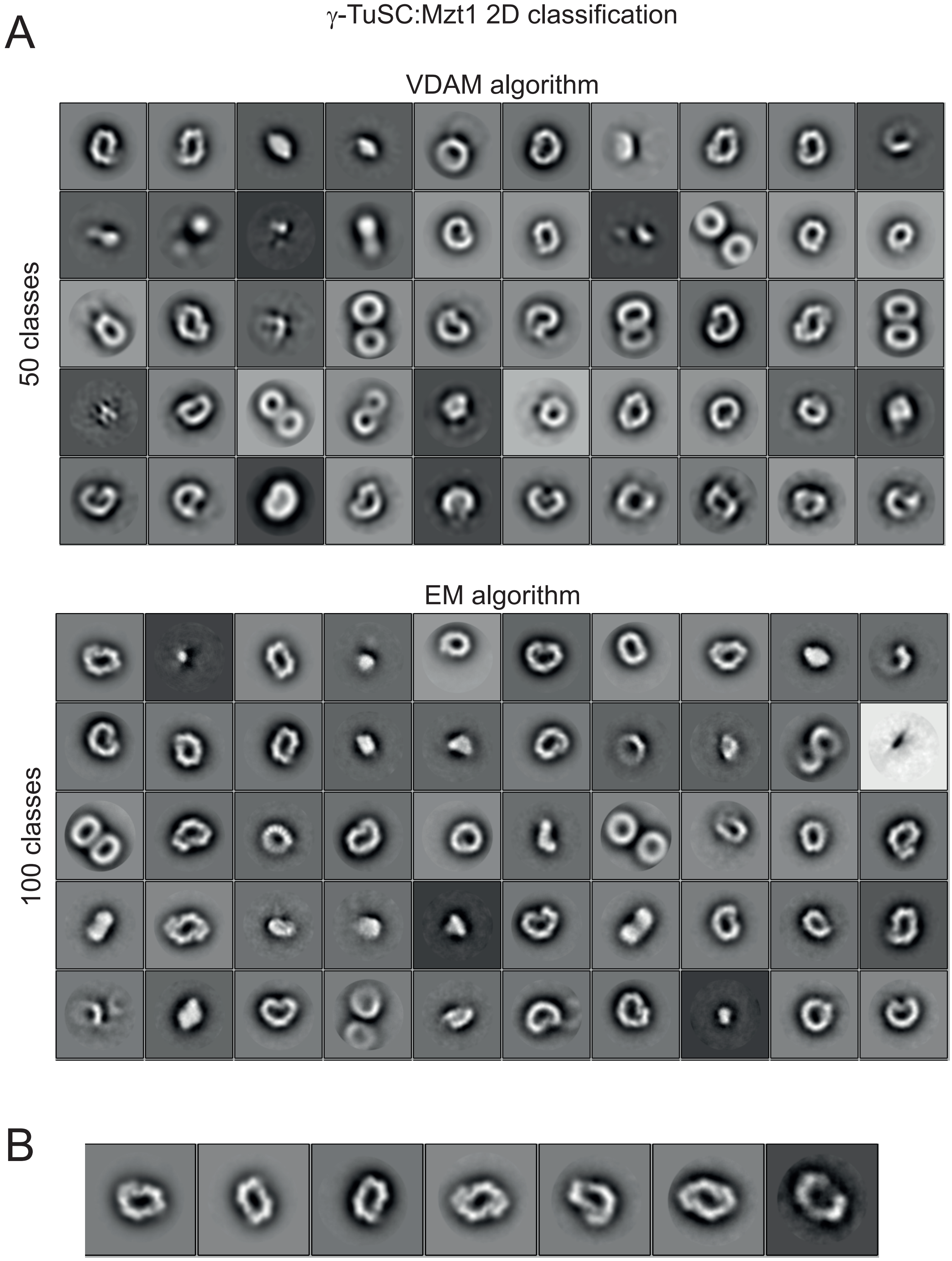


**Supplementary Figure 8: γ-TuSC:Mzt1 2D classification.**
**A**) 2D classifications using the VDAM algorithm (50 classes specified) or EM algorithm (100 classes specified). For the EM classification, only the top 50 most populated classes are shown. Classes are ordered by the number of particles contributing to each class, with the top-left class containing the most particles, and then row-by-row, with the bottom-right class containing the fewest particles. **B**) 2D classes from γ-TuSC:Mzt1 EM algorithm dataset that have two-fold symmetry (several are also shown in A).

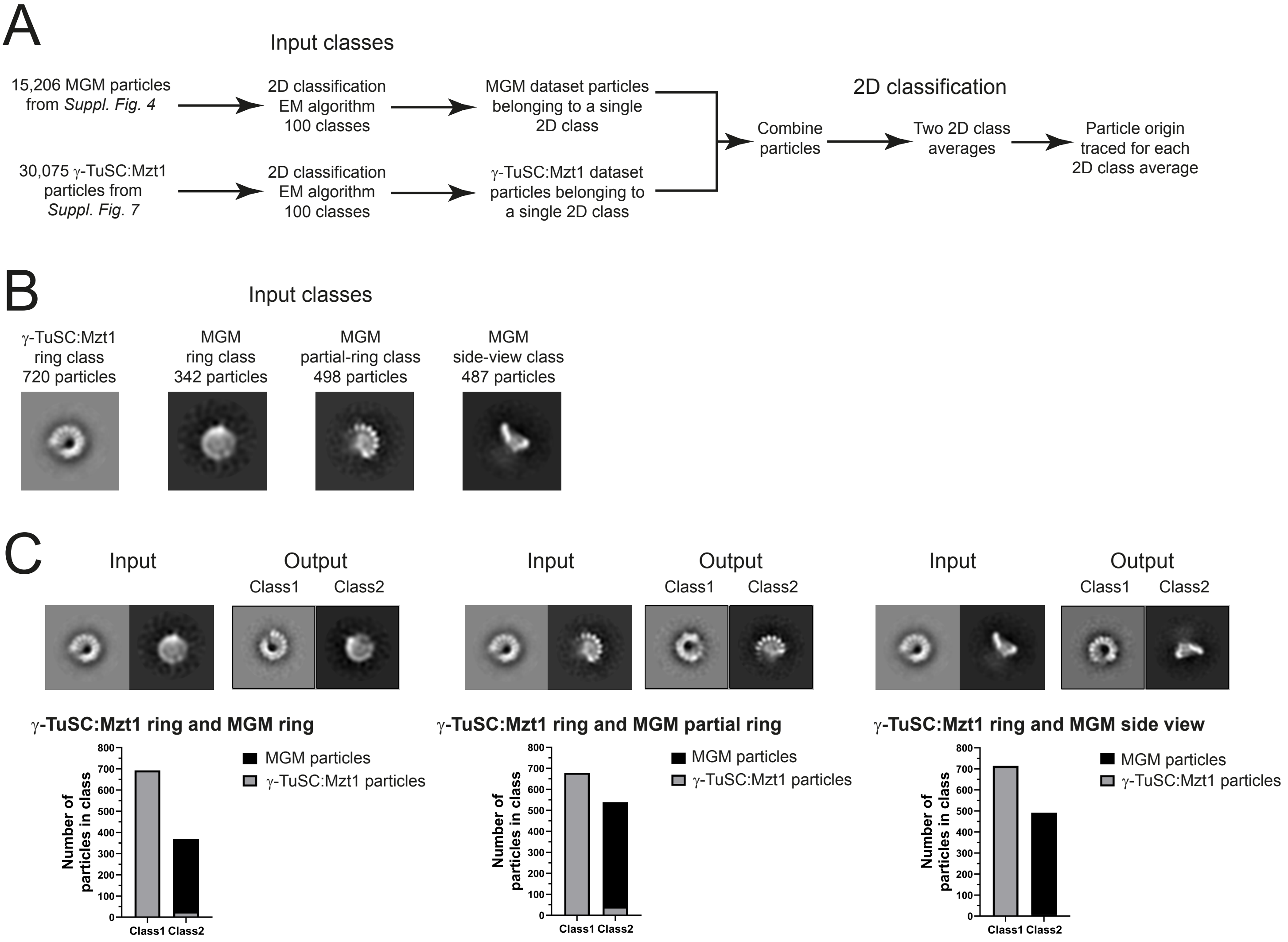


**Supplementary Figure 9: Comparison of γ-TuSC:Mzt1 and MGM conformations by 2D classification.** **A**) Schematic of analyses. Prior processing of inputs is shown in Suppl. Figs 4 and 7; arrows on those figures show the point at which data were used for these analyses. **B**) The four input classes used for pairwise analyses. γ-TuSC:Mzt1 ring class was compared pairwise with three individual MGM classes. **C**) Input classes and output classes for each of the three pairwise analyses. Bar charts show the number of MGM and γ-TuSC:Mzt1 particles present in each output class (Class1, Class2).

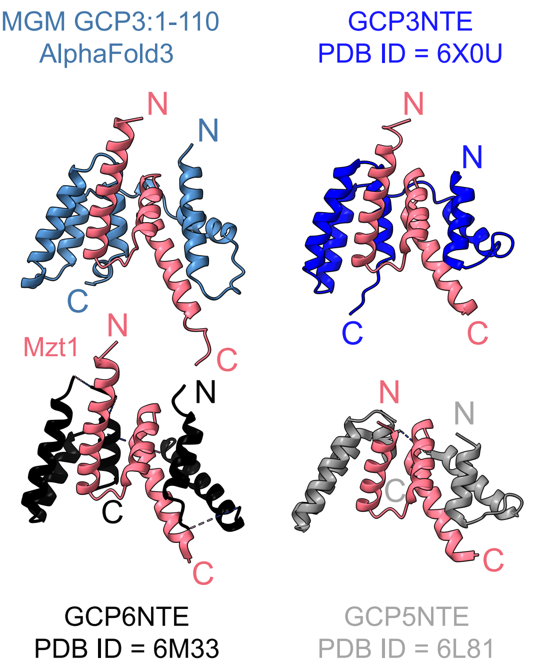


| Previously determined structure: | Cα RMSD compared to GCP3NTE-Mzt1 model (Å): |
|---|---|
| Mzt1-GCP3NTE | 1.05 |
| Mzt1-GCP6NTE | 0.97 |
| Mzt1-GCP5NTE | 0.66 |

**Supplementary Figure 10: Comparison of the MGM GCP3NTE-Mzt1 AlphaFold3-predicted structure with *H. sapiens* GCPNTE-Mzt1 structures previously determined by single particle analysis or X-ray crystallography.**
Colors of labels match the corresponding polypeptides. Residues nearest to N- and C-terminal residues (according to primary sequence) are labelled “N” and “C”, respectively

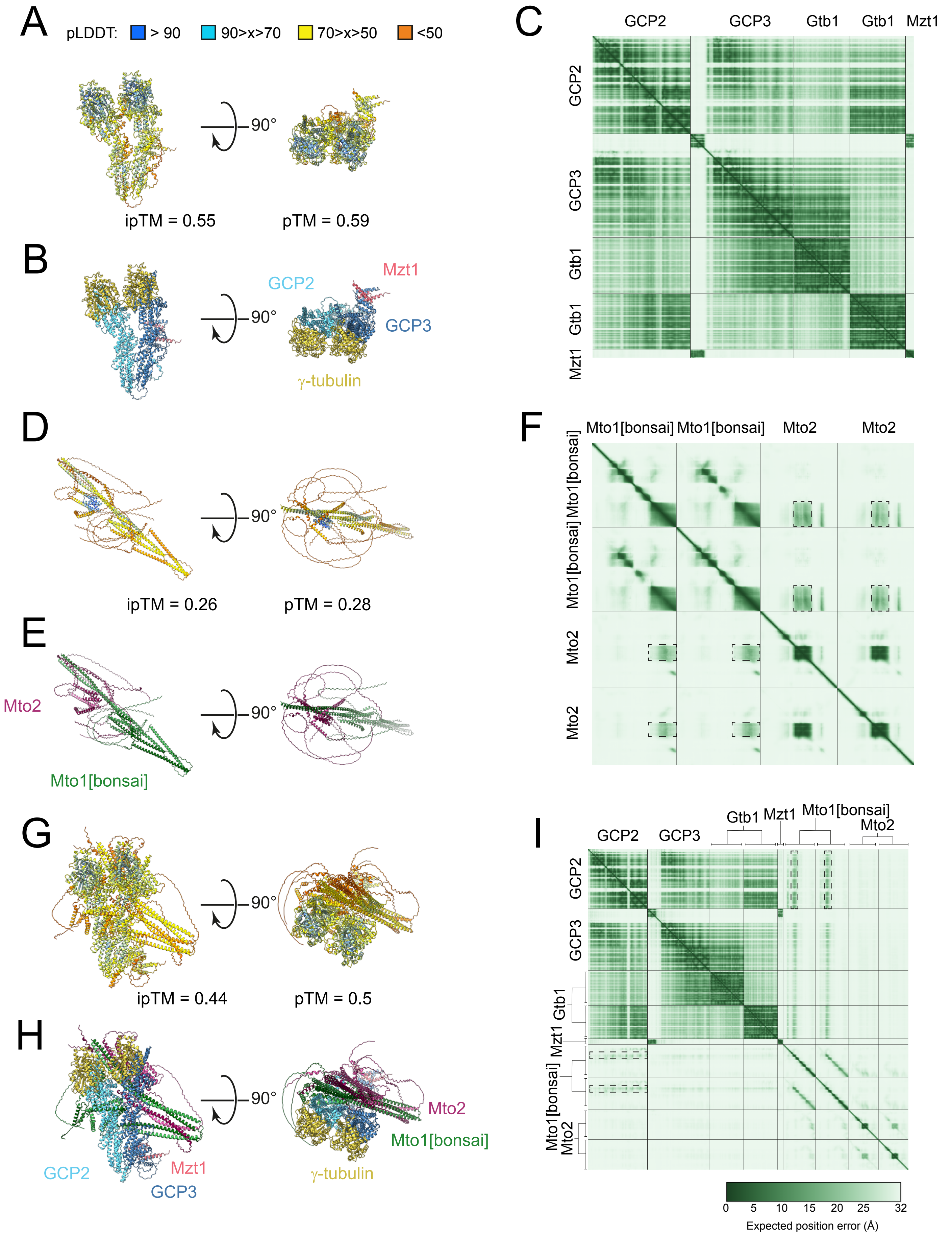


**Supplementary Figure 11: AlphaFold3 models and model statistics for different MGM subcomplexes.** **A-C**) γ-TuSC:Mzt1 heteropentamer. **D-F**) Mto1[bonsai]:Mto2 heterotetramer (2:2). **G-I**) MGM heterononamer subcomplex. Panels A, D, and G are colored by predicted local distance difference test (pLDDT) values. Panels B, E, and H are colored by protein identity. Panels C, F, and I show predicted aligned error (PAE) plots. For each model, interface predicted template modeling (ipTM) and predicted template modeling (pTM) scores are also shown. Dashed boxes on PAE plot in F indicate regions involved in interaction between Mto2 twin cysteine domain (Mto2-TCD) homodimer and Mto1[bonsai] coiled-coil dimer. Dashed boxes on PAE plot in I indicate regions involved in interaction between Mto1[bonsai] CM1 domains and GCP2.

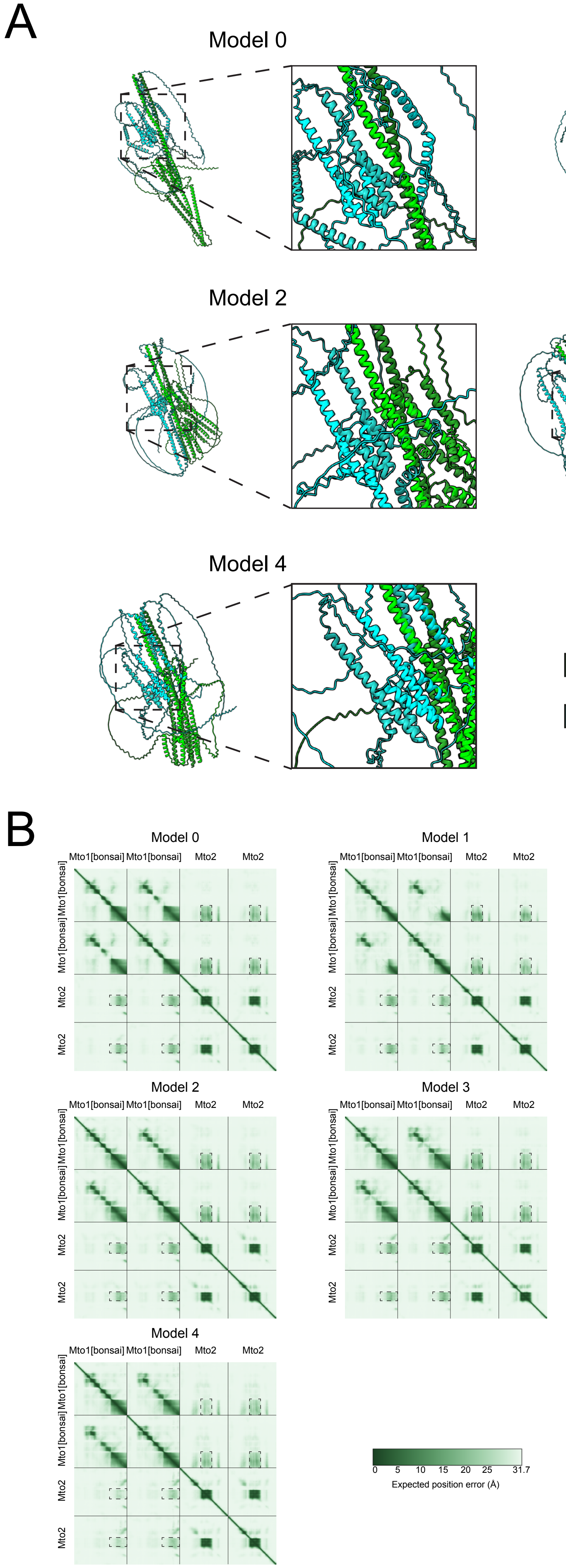


**Supplementary Figure 12: AlphaFold3 models of Mto1[bonsai]:Mto2 heterotetramer (2:2).** **A**) Cartoon representations of the five models (Model 0 to Model 4). **B**) Predicted aligned error (PAE) plots for each model. Interface predicted template modeling (ipTM) scores for Models 0-4 were 0.26, 0.25, 0.26, 0.27, and 0.26, respectively. Predicted template modeling (pTM) scores for Models 0-4 were 0.28, 0.27, 0.28, 0.29, and 0.28, respectively. Dashed boxes (zoomed) in A indicate interaction between the Mto2 twin cysteine domain (Mto2-TCD) homodimer and the Mto1[bonsai] coiled-coil dimer, which is present in all models. Dashed boxes in B indicate regions of Mto1[bonsai] and Mto2 involved in the interaction.

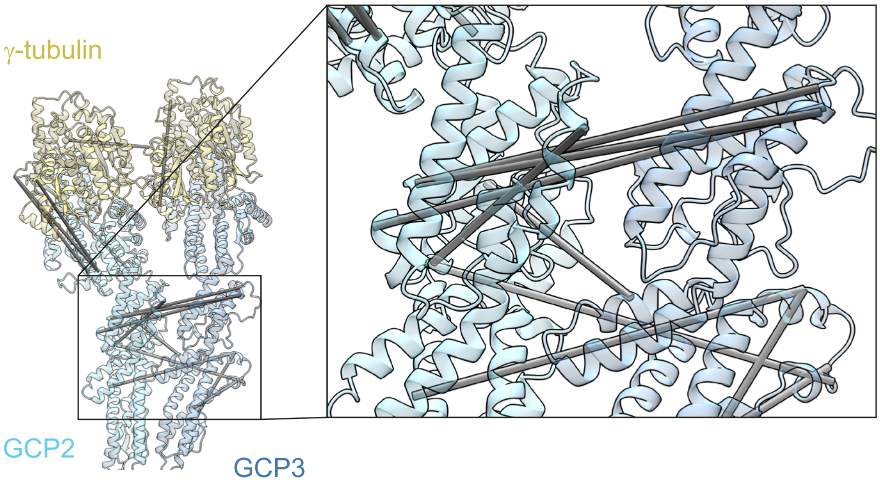


**Supplementary Figure 13: Crosslinks involving γ-TuSC that violate the 30 Å distance constraint when mapped to a single γ-TuSC.** AlphaFold3 predicted model for γ-TuSC, with unstructured segments removed, shown with experimentally determined chemical crosslinks (gray rods) having Cα - Cα distances >30 Å within the context of the model. Many of these violations can be explained by inter-γ-TuSC interactions (see “one-and-a-half” γ-TuSC model in Figure 4). The different shading of gray rods is a consequence of the angles of illumination used for displaying the model.

**Supplementary Table 1:** Table of cross-correlation values for comparing open and closed $\gamma$-TuRCs to *S. pombe* 10 $\gamma$-tubulin MGM, 14 $\gamma$-tubulin MGM and single ring $\gamma$-TuSC:Mzt1 reconstructions.

| | $\gamma$-TuRC Sample | 10 $\gamma$-tubulin MGM reconstruction | 14 $\gamma$-tubulin MGM reconstruction | $\gamma$-TuSC:Mzt1 reconstruction |
|---|---|---|---|---|
| Open conformations | Native *X. laevis* PDB ID = 6TF9 | 0.846 | 0.797 | 0.823 |
| | Recombinant *H. sapiens* PDB ID = 7QJ5 | 0.852 | 0.793 | 0.797 |
| | Native *H. sapiens* PDB ID = 6V6S | 0.853 | 0.799 | 0.821 |
| | Recombinant *H. sapiens* PDB ID = 7AS4 | 0.855 | 0.797 | 0.821 |
| Closed conformations | Recombinant *H. sapiens* microtubule-bound $\gamma$-TuRC PDD ID = 8VRK | 0.843 | 0.802 | 0.819 |
| | Native spindle pole body *S. cerevisiae* microtubule -bound $\gamma$-TuRC PDB ID = 8QV2 | 0.867 | 0.806 | 0.840 |
| | Recombinant *H. sapiens* microtubule-bound fully closed $\gamma$-TuRC | 0.843 | 0.801 | 0.839 |
